# A quorum of mechano-sensing fungal consortia

**DOI:** 10.1101/2022.10.24.513463

**Authors:** M. García-Navarrete, D. Ruiz Sanchis, I. Sánchez-Muñoz, S. González-Ruiz, M. Avdovic, S. Atienza-Sanz, K. Wabnik

## Abstract

Bacteria use Quorum sensing (QS) to coordinate gene expression in dense cell populations. Here, we show that eukaryote *S. Cerevisiae* uses an alternative strategy, the quorum mechano-sensing (QMS), to resolve growth conflicts in the fungal consortia. QMS connects the biomechanical signal perception through adhesin FLO11 and transmembrane histidine kinase SLN1, triggering an intracellular signaling cascade for the cell density-dependent regulation of gene expression. Both cis and trans interactions of FLO11 are required for the inhibition of SLN1 and involve the extracellular fibronectin type III-like domain of FLO11. Genetic deletion of FLO11 removes inhibition of SLN1, associated with the spontaneous activation of gene expression whereas overproduction of FLO11 strengthens the inhibitory effect of FLO11 on SLN1. Therefore, adjusting the amount of FLO11 directly scales with the level of SLN1 inhibition, forecasting the outcome of growth competition at the macroscopic scale. Furthermore, the integration of an orthogonal synthetic circuit downstream of SLN1 allows for QMS-controlled regulation of gene expression in cell populations. Our study reveals a molecular pathway connecting FLO11 adhesion to SLN1-dependent intracellular regulation of gene expression in fungi. FLO11 and SLN1 coordinate kin recognition and growth conflict resolution through gene expression in dense fungal populations. This study challenges the classical view of chemically-driven QS and provides new strategies for controlling population growth through quorum mechano-sensing.

## Main

Quorum sensing (QS) allows bacteria to communicate chemical messages across entire populations to recognize familiar kin^1^. Actively growing microbial cells produce diffusing QS signals at a critical concentration threshold that directly scales with population density^1^. This threshold triggers synchronized gene expression across the microbial consortia, providing a mechanism for cellular noise buffering and robust and coordinated colony-level responses^2,3^. Similar QS-like mechanisms may exist in eukaryotes such as fungi but underlying molecular mechanisms remain an enigma^4,5^.

It is well established that the kin recognition in unicellular fungi such as yeast *S. Cerevisiae* requires intercellular(trans) and in the same cell (cis) mechanical interactions mediated by the FLO11 cell wall-resident adhesin^6–10^. Again, how FLO11-mediated cell contacts are mechanistically interpreted by cells to trigger Kin recognition is however unclear. One possibility is that FLO11 requires a partner to transmit a signal to the cell interior to control gene expression required for kin recognition and growth competition in mixed consortia. Nevertheless, no such partner proteins were identified to date. Mechanical stresses such as osmotic conditions and cell wall integrity are sensed by transmembrane histidine kinase SLN1^11–17^ that transmits a signal via a multistep phosphorelay cascade to control gene regulation^18–20^. The molecular basis of SLN1-mediated mechanical stress perception in the cell wall remains still unknown. Intriguingly, the SLN1 activity can be modulated by cell-wall resident mannoproteins^21^.

Herein, we hypothesize that SLN1 could be so far unknown partner for FLO11 that connect extracellular Kin recognition via contact-based mechanics with intracellular signaling controlling downstream gene expression.

## Results

### Kin detection in fungal consortia requires the new SLN1-dependent intracellular signaling route

To test this idea, we used two strains of yeast *S. Cerevisiae* with different competence for FLO11 expression to establish a mixed fungal consortium model (Fig. 1A). CEN.PK2 strain(‘Kin’) has lost major flocculation genes during the evolution^22^ but retained FLO11 which is the central regulator of biofilm formation and invasive growth^8,23–26^. In contrast, the BY4741 strain(‘Intruder’) is a descendent of wild-type yeast S288C that does not express FLO11 due to deleterious mutations in FLO8 transcriptional activator^6,9^. Both strains show similar growth kinetics when cultivated alone (Fig. S1), but only ‘Kin’ is competent to express FLO11 (Fig. 1A). We next checked whether the SLN1-dependent pathway (Fig. 1B) in the ‘Kin’ is active in this model consortia using the short-lived NeonGreen reporter (dNeon) placed under the control of the OCH1 promoter^27^ or its engineered synthetic variant^28^. The activity of both promoters presumably depends on transcription factor SKN7 which is activated by SLN1 through trans-phosphorylation^20,27^ (Fig. 1B). In either scenario, the ‘Kin’ reporter was, however, inactive when cocultured with the ‘Intruder’ strain (Fig. S2A).

**Fig. 1.**
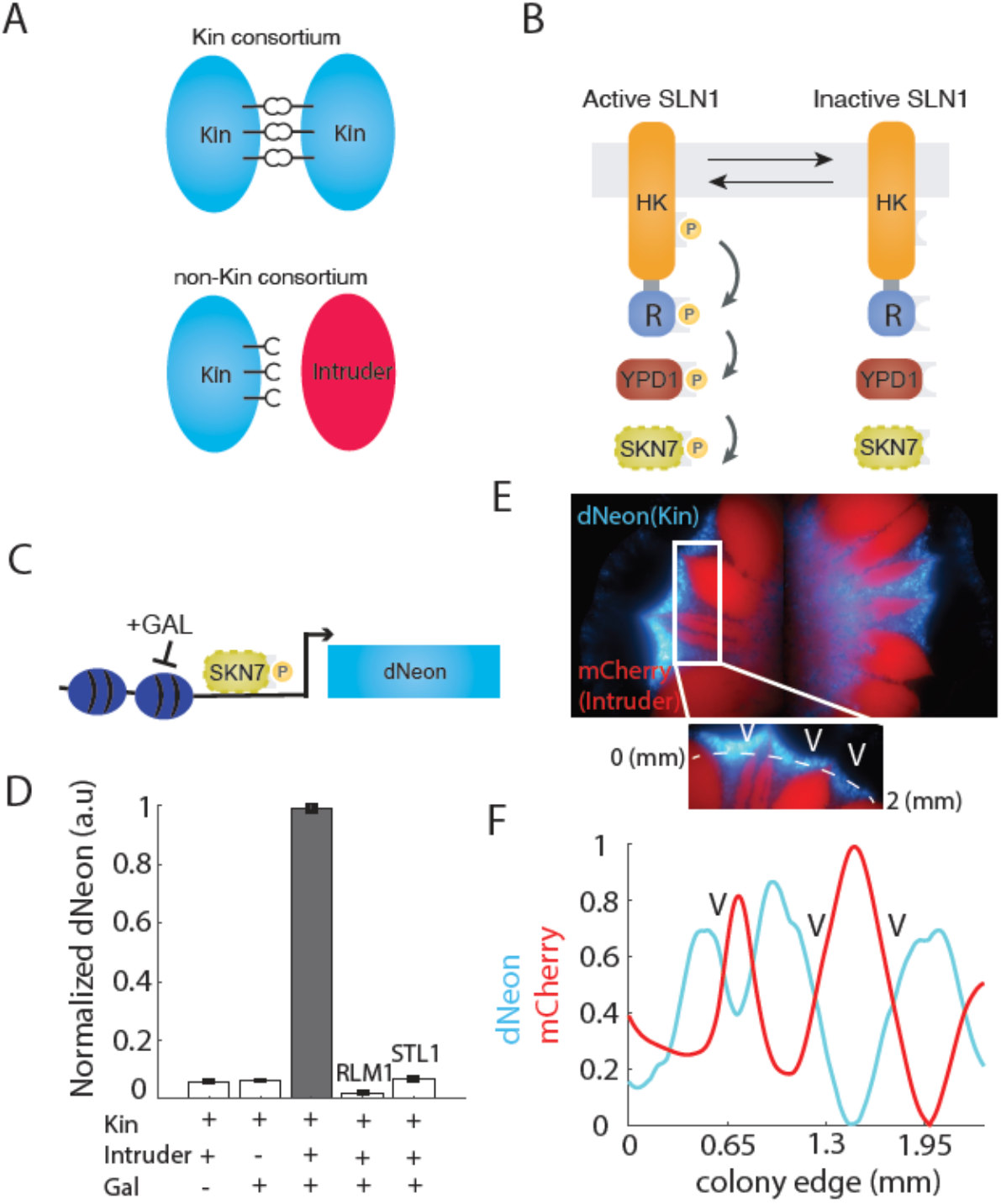
Contact-based detection mechanism resolves growth conflicts in yeast consortia. **A,** Schematic of kin and non-kin strain consortia. ‘Kin’ (CEN.PK2) and ‘Intruder’ (BY4741) are haploid yeast strains. ‘Kin’ can express FLO11 whereas ‘Intruder’ has no expression of FLO11 due to a FLO8 transcriptional activator defect. **B**, Schematics of the canonical SLN1-dependent signaling cascade. SLN1 in an active phosphorylated state transfer the phosphoryl group from the receiver domain (R) to YPD1 and subsequently onto the SKN7 transcription factor. **C**, Basic schematic of engineered SKN7-responsive promoter. Galactose promotes chromatin opening followed by active SKN7 attachment to DNA. This promoter drives dNeon reporter to monitor dynamic changes in fluorescence signal. **D**, Normalized dNeon activation in different co-culturing conditions. Note that the observed response is not triggered by wall stress (pRML1 reporter^13^) or changes in osmotic conditions (pSTL1 reporter^12^). Note that only the presence of both galactose and ‘Intruder’ triggers dNeon activation. **E**, Example growth patterns of mixed consortia (1:1 initial inoculation at OD_600_=0.1 each) show astonishing growth arrangements with ‘Kin’ protruding from the colony peripheral after 5-7 days of growth in standard conditions on agar plates. ‘Kin’ dNeon reporter is shown in cyan and ‘Intruder’ expressing constitutively mCherry is shown in red. **F**, Representative trend of growth pattern on the colony peripheral (n=10), arc on colony periphery(edge) was depicted by the dashed white curve. Note the generation of a flower-like pattern with ‘Kin’ growth protrusions.

Previous studies suggest that SKN7 may require co-activators and does not act as the sole transcriptional regulator^29–31^. We sought that SKN7 may require a coactivator to strongly induce gene expression following its activation by SLN1. To investigate this possibility, we engineered a bidirectional synthetic promoter (denoted as pSynSKN7) to facilitate the galactose-promoted chromatin opening^32^ and subsequent recruitment of activated SKN7 to control pSynSKN7 promoter activity (Fig. 1C, Fig. S2B). Remarkably, we reported that the dNeon reporter in the ‘Kin’ driven by pSynSKN7 promoter was activated solely in the presence of ‘Intruder’ and galactose, whereas it remained inactive under all control conditions (Fig. 1D). This result suggests that active SKN7 as well as the difference in the competence for FLO11 between ‘Kin’ and ‘Intruder’ may play an important role in the dNeon reporter activation because ‘Kin’-‘Kin’ cocultures show no detectable reporter activity in the presence of galactose (Fig. 1D). Then, we asked whether this activation requires canonical HOG1-dependent osmotic response downstream of SLN1^12^ or could be triggered by cell wall stress^13^. We could not detect any reporter activity that was driven from commonly used hyperosmotic^12^ and cell wall stress responsive^13^ promoters in co-cultures of ‘Kin’ and ‘Intruder’(Fig. 1D), These data suggest that neither hyperosmotic response nor cell wall stress seems to underly observed activation of dNeon reporter in ‘Kin’. Based on these findings we concluded that a previously unknown SLN1-dependent signaling route could be involved.

### Dynamic, cell density-dependent mechanism connects mechanical cell contacts with activity of SLN1 intracellular pathway

We then inspected if there are morphogenetic changes associated with dNeon activation in the growth competition between ‘Kin’ and ‘Intruder’. We co-cultured Kin’ and ‘Intruder’ over 5 to 6 days in normal conditions on solid agar plates. Remarkably, we recorded the emergence of flower-like patterns that were dominated by protruding ‘Kin’ populations (Fig. 1E). These growth patterns of ‘Kin’ (cyan) generated a sharp boundary interface with the ‘Intruder’ (red) on the consortium periphery (Fig. 1F and Fig. S2C). By creating these tight barriers, ‘Kin’ was able to eventually outcompete the ‘Intruder’ (Fig. 1E).

To understand better the kinetics of this ‘Kin’ behavior, we performed time-course experiments to track dNeon reporter dynamics over time in uniformly mixed liquid cocultures of ‘Kin’ and ‘Intruder’. Unexpectedly, we found that the dNeon reporter was rapidly activated at the specific cell density threshold (THR) represented as OD_600_ measured at 20% of the peak reporter fluorescence. This THR corresponds to the late-exponential phase/early stationary phase (OD_600_ ~ 1.15, Fig. 2A). The dNeon reporter response was dramatically increased following further increase of cell densities (Fig. 2A), demonstrating characteristics of the QS-like phenomena (Fig. 2B). Because chemical compounds, such as 2-phenylethanol, tryptophol, and tyrosol^33,34^ produced by yeast have been suggested to possibly act as QS molecules, we investigated whether any of these chemicals trigger dNeon reporter. We culture ‘Kin’ with concentrated extracts of media in which ‘Intruder’ was grown that would be enriched in putative QS-like molecules. Nevertheless, dNeon reporter was inactive under these conditions (Fig. S3), again indicating that the trigger is not chemical in nature.

**Fig. 2.**
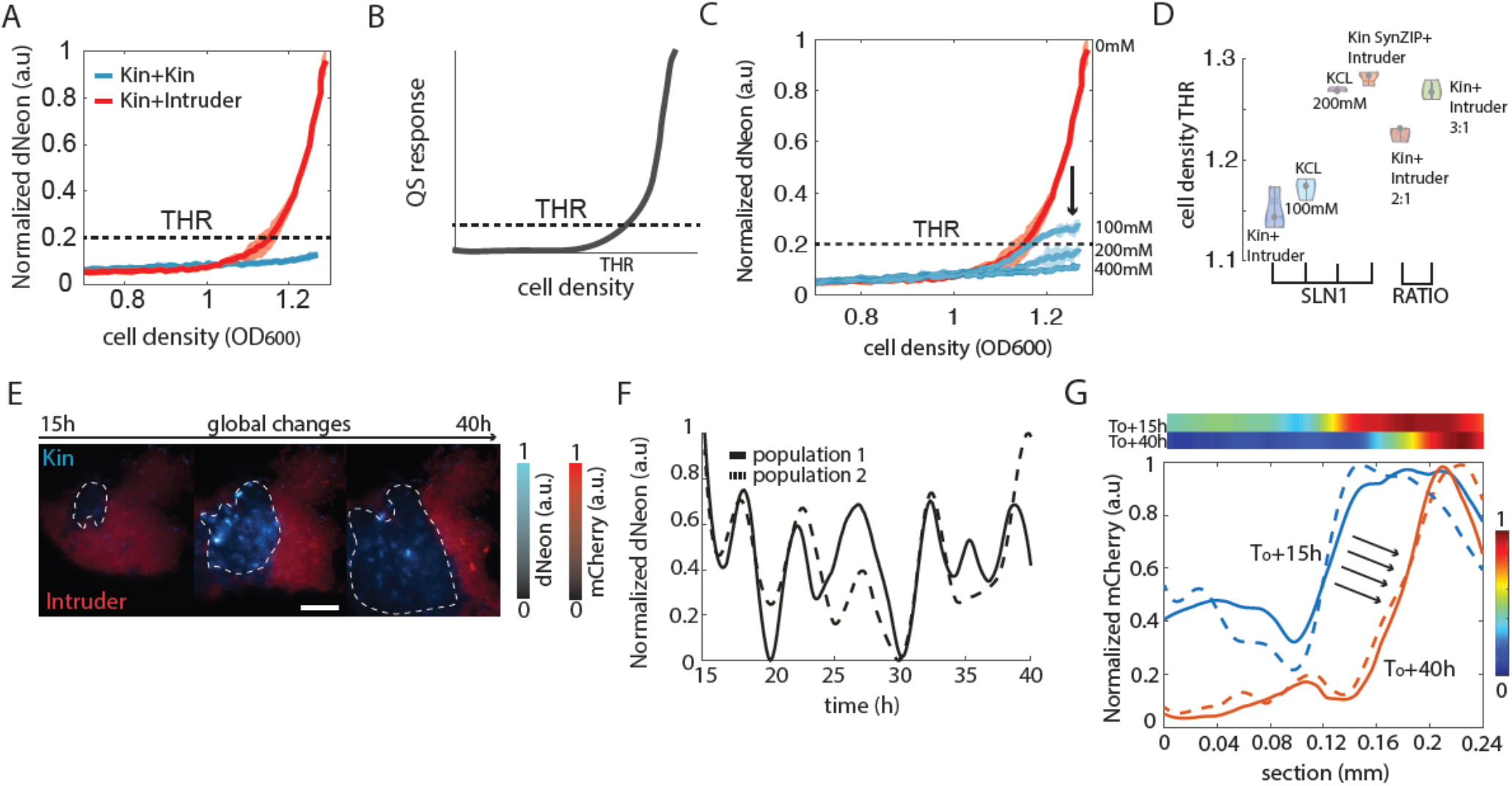
Quorum mechano-sensing(QMS) controls dynamic population-level gene expression response of Kin to the presence of Intruder. **A,** Time-lapse kinetics of ‘Kin’ activation (QMS curve) in the presence (red curve) or absence of ‘Intruder’ (blue curve) (A). Note that cell density threshold (THR) for dNeon reporter activation corresponds to early stationary phase of cell growth (OD_600_ > 1) at the 20% of maximal reponse. **B**, The model kinetics of classical QS response. **C**, The effect of hyperosmotic stress in the Kin-Intruder cocultures. Note progressive shift of THR to higher OD_600_ and gradual decrease in the amplitude of response. **D**, Changes in THR values by manipulating the SLN1-signalling pathway (chemically or genetically) or increasing the initial inoculation of Kin (increasing FLO11 presence). **E**, Snapshots from time lapse experiments of mixed ‘Kin’ and ‘Intruder’ (1:1) consortia performed in the microfluidic device. dNeon fluorescence (’Kin’) is shown in cyan and ‘Intruder’ is marker in red with constitutive mCherry reporter. Note transient contacts between strains trigger dNeon response following the switch-like dynamics. **F**, Representative time-course plots (n=10) from microfluidic experiments presented in (E). Note the transient ON-OFF dynamics of ‘Kin’ in response to the presence of ‘Intruder’. **G**, Time evolution of growth conflict resolution between ‘Kin’ and ‘Intruder’ in microfluidic environment (two examples of trap sections are shown as solid and dash curves, respectively). The ‘Intruder’ presence shown with mCherry reporter was quantified over time. In ~24h (red curves) ‘Intruder’ occupancy drastically decreases to nearly 50% of its original size (blue curves). This is a consequence of aggressive growth of the ‘Kin’. QMS curves show mean trends with 95% confidence intervals collected from three independent replicates.

We then tested whether the ‘Kin’ response can be attenuated by imposing the hyperosmotic stress with KCL which is known to deactivate SLN1^12^. Indeed, treatments with salts led to a gradual decrease of dNeon reporter response and a shift of THR towards higher values (Fig. 2C and D), further suggesting that the activation of ‘Kin’ relies on a phosphorylated(active) form of SLN1. Similarly, the hyperosmotic stress can lead to the tight clustering of SLN1 and its subsequent deactivation^18^. To further test whether imposing dimerization of SLN1 could impair dNeon expression by deactivating SLN1 we constructed a chimeric SLN1 in which we swapped the extracellular domain for complementary synthetic leucine zippers^35^ (Fig. S4A). Indeed, this synthetic pair of chimeric SLN1s decreased the activity of SLN1 and attenuated dNeon reporter expression (Fig. S4B).

Next, to quantify the dynamics of dNeon activity in ‘Kin’, we cocultured both strains in the microfluidic device (Fig. S5) under continuous media perfusion in the controlled environment. Surprisingly, the experiments on a chip revealed transient switch-like activation of the dNeon reporter following the spontaneous establishment of contacts with ‘Intruder’ (Fig. 2E, F, Fig. S6A, and Video S1). Furthermore, the presence of dNeon fluorescence preceded ‘Kin’ aggregation, creating a stiff growth barrier to outcompete the “Intruder” (Fig. 2G, Fig. S6B) similar to that observed in co-cultures grown on solid media (Fig. 1E). Thus, a response to a mechanical contact between ‘Kin’ and ‘Intruder’ exhibits a characteristic of QS, we refer to it onwards as Quorum mechano-sensing (QMS).

Our experiments indicate that dNeon expression was inhibited until a specific cell density was reached. This suggests there may be a general inhibitor of the SLN1 pathway that controls Kin recognition. We then sought to test if increasing the initial amount of Kin in cocultures with ‘Intruder’ would increase the time of dNeon inhibition by shifting THR to higher values. Indeed, 2:1 and 3:1 ratios of Kin to Intruder led to longer inhibition of the reporter and increased THR (Figs. 2D, and S7A, B), suggesting that the inhibition originates from ‘Kin’. These data indicate that the ‘Kin’ strain behaves as a density-dependent trigger in the presence of ‘Intruder’.

### FLO11 adhesin controls the timing and steepness of QMS response

Since there is a clear difference in the genetic competence for FLO11 expression between ‘Kin’ and ‘Intruder’, we tested whether the observed dynamic dNeon response in ‘Kin’ relies on the FLO11 activity. We used CRISPR-Cas to knockout Flo11 in ‘Kin’, to remove FLO11. Remarkably, flo11-deficient ‘Kin Δflo11’ strain was strongly activated even in the absence of ‘Intruder’ (Fig. 3A). Also, we noticed early activation and lack of inhibition as compared to cocultures of ‘Kin’-‘Intruder’ in which FLO11 was intact (Fig. 3A, G). Next, we cocultured ‘Kin Δflo11’ with ‘Kin’ to test whether FLO11-dependent inhibition of dNeon reporter can be partially restored. Our experiment confirmed that FLO11 presence in contacts with Kin lacking FLO11 can partially restore this inhibition as seen by increased THR (Fig. 3A, G). Our data confirm the inhibitory role of FLO11 in the dNeon reporter activation. Furthermore, it is plausible that FLO11 could directly control THR and the steepness of QMS response.

**Fig. 3.**
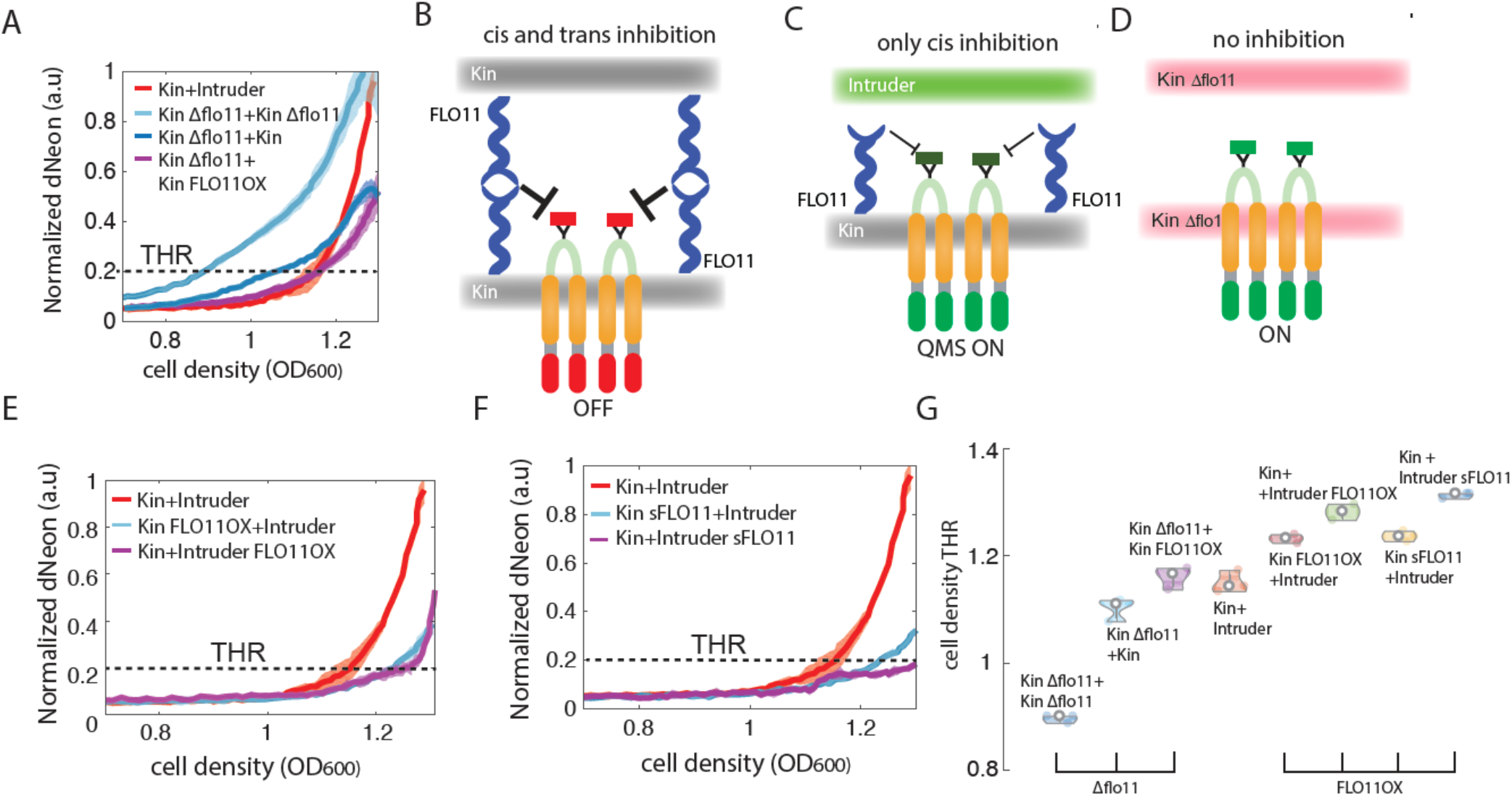
FLO11 promotes SLN1 inhibition and modulates QMS responsiveness characteristics. **A,** QMS curves in wt and flo11 mutant of ‘Kin’ (Kin Δflo11). The deletion of FLO11 in ‘Kin’ shows no repression and early dNeon signal accumulation. Note that coculturing with Kin or FLO11 overexpressing strain restores the inhibition of dNeon reporter expression to the level similar to that of Kin-Intruder cocultures. **B-D,** Schematic of the QMS mechanism. In control condition FLO11 in isogenic ‘Kin’ cocultures creates stable trans and cis contacts, completely repressing the SLN1 activation(B). Intruders lack FLO11 during contacts with Kin and only cis contacts are permitted, yielding non-complete repression (C). The complete lack of FLO11 in isogenic Kin Δflo11 cultures results in strong activation of dNeon reporter, turning off the QMS mechanism. **E, F**, Coculturing experiments with strains overexpressing FLO11 full length(E) or sFLO11 genes(F). Note that boosting FLO11 in ‘Kin’ has opposite effect to its deletion (A). Increase in FLO11 in ‘Kin’ compared to ‘Intruder’ resulted in stronger inhibition of dNeon response. Increasing contacts with homotypic FLO11 interactions through explicitly FLO11 extracellular domain in ‘Intruder’ boost inhibition of dNeon expression. This effect was prominent in chimeric protein sFLO11 that contains N-terminus domain of FLO11. **G**, THR changes in different cocultures lacking or overexpressing FLO11 adhesin. QMS curves show mean trends with 95% confidence intervals collected from three independent replicates.

Based on these findings we propose the following model:

1. the presence of FLO11 inhibits the SLN1 pathway as seen for ‘Kin’-‘Kin’ cultures possibly through direct or indirect cis and trans interaction with the SLN1 ectodomain(Fig. 3B). (2) increased cell densities in ‘Kin’-‘Intruder’ consortia could increase the probability for transient ‘Kin’-‘Intruder’ contacts that would partially relieve the SLN1 pathway from FLO11-dependent inhibition (Fig. 3C). (3) the complete removal of FLO11 would allow for early activation of the SLN1 pathway (Fig. 3D).

According to this model, increasing FLO11 expression in ‘Kin’ should further inhibit reporter expression and delay response. To test this concept, we generated the ‘Kin’ strain (‘Kin FLO11OX’) by overexpressing FLO11. Notably, we found that coculturing ‘Kin FLO11OX’ with ‘Intruder’ resulted in a 50% weaker response compared to controls (Fig. 3E). Furthermore, the dNeon activation was delayed as demonstrated by increased THR value (Fig. 3E, G). Similarly, producing FLO11 in ‘Intruder’ showed delayed reporter response due to the longer inhibition of activation as manifested by high THR values (Fig. 3E, G). Strikingly coculturing ‘Kin Δflo11’ with ‘Kin FLO11OX’ restored the inhibition of the dNeon reporter to a level similar to that of ‘Kin’-‘Intruder’(Fig. 3A, G). Thus, all these data are in agreement with our model (Fig. 3B-D). We then asked which part of FLO11 would be involved in the inhibition of SLN1 in ‘Kin’. For that purpose, we constructed a synthetic membrane receptor spanning cell wall by fusing nano spring-like ectodomains of WSC1 and MID2 cell-wall proteins^36,37^ with the adhesin N-terminal part of FLO11 mediating cis- and trans-interactions^38–42^ (Fig. S8). The application of nano springs allows the mechanical unfolding of chimera in the cell wall during FLO11-mediated contacts. We expressed this chimeric protein in either ‘Kin sFLO11 ‘ or ‘Intruder sFLO11 ‘ strains and monitor dNeon reporter activity in ‘Kin’. Intriguingly, the induced expression of the chimeric membrane-bound FLO11 receptor showed consistent but much more noticeable changes in dNeon reporter activity as compared to FLO11 full-length overexpression (Fig. 3F, G). Notably, we reported a strong inhibition of reporter signal in ‘Kin sFLO11’ in coculture with ‘Intruder’ (Fig. 3F), similarly cocultures of ‘Kin’ with ‘Intruder sFLO11 ‘, resulted in much more delayed reporter activation (Fig. 3F, G).

Finally, we asked whether the same mechanism would apply to ‘Intruder’ which normally lacks FLO11 due to FLO8 mutation^6,9^. Indeed, coculturing ‘Intruder dNeon’ with ‘Intruder’, led to the spontaneous activation of dNeon fluorescence as observed in the case of ‘Kin Δflo11’ (Fig. S9A). However, combining ‘Intruder with dNeon’ with ‘Kin’ restored QMS mechanisms by shifting THR to higher cell densities (Fig. S9A, B). Finally, we overexpressed FLO11 in ‘Intruder’ or ‘Intruder dNeon’ simultaneously monitoring reporter activity in cocultures. We found that increased levels of FLO11 correlated with higher inhibition of dNeon reporter signal in both cis and trans configurations (Fig. S9A, B), similar to that observed in ‘Kin’. All these data indicate that modulating FLO11 in the consortia allows control of SLN1-dependent gene expression through the QMS mechanism. Both THR and amplitude of colony-level response can be fine-tuned through the SLN1 pathway by changing the relative ratio of adhesin expression in competing strains.

### Connecting QMS to a synthetic circuit for orthogonal control of gene expression in dense cell populations

Our findings indicate that the QMS mechanism mediates SLN1 activation through FLO11-dependent cell contacts. QMS leads to a subsequent two-step phosphorylation cascade involving phosphor relay onto the SKN7 transcription factor^12–20^ whose activity is quantified by dNeon reporter. Since SLN1 mutants are lethal and mutations in downstream components such as SKN7 could produce unwanted pleiotropic effects, we sought to further explore the pathway downstream of SLN1 by constructing a synthetic orthogonal pathway to plug QMS to downstream regulation of gene expression. Bacterial two-step HK kinase systems^43–45^ offer an excellent opportunity to introduce heterologous HK-based regulation into fungi because of evolutionary similarities between fungi and bacterial HK systems. We chose a specific FixL-FixJ system from the nitrogen-fixing bacteria *Rhizobium* meliloti^46,47^ as this system has a lower likelihood of being activated by the yeast YPD1 kinase (Fig. 4A, Fig. S10A). To assemble a synthetic rerouting, we created a chimeric HK by fusing the signal peptide, transmembrane, and HAMP domains of SLN1 with the HK domain of FixL HK (Fig. S10A). FixJ transcriptional regulator was fused to the transactivation domain from Herpes virus (VP64)^48^. We then placed our dNeon reporter under the control of synthetic FixJ-regulated promoter^49,50^.

**Fig. 4.**
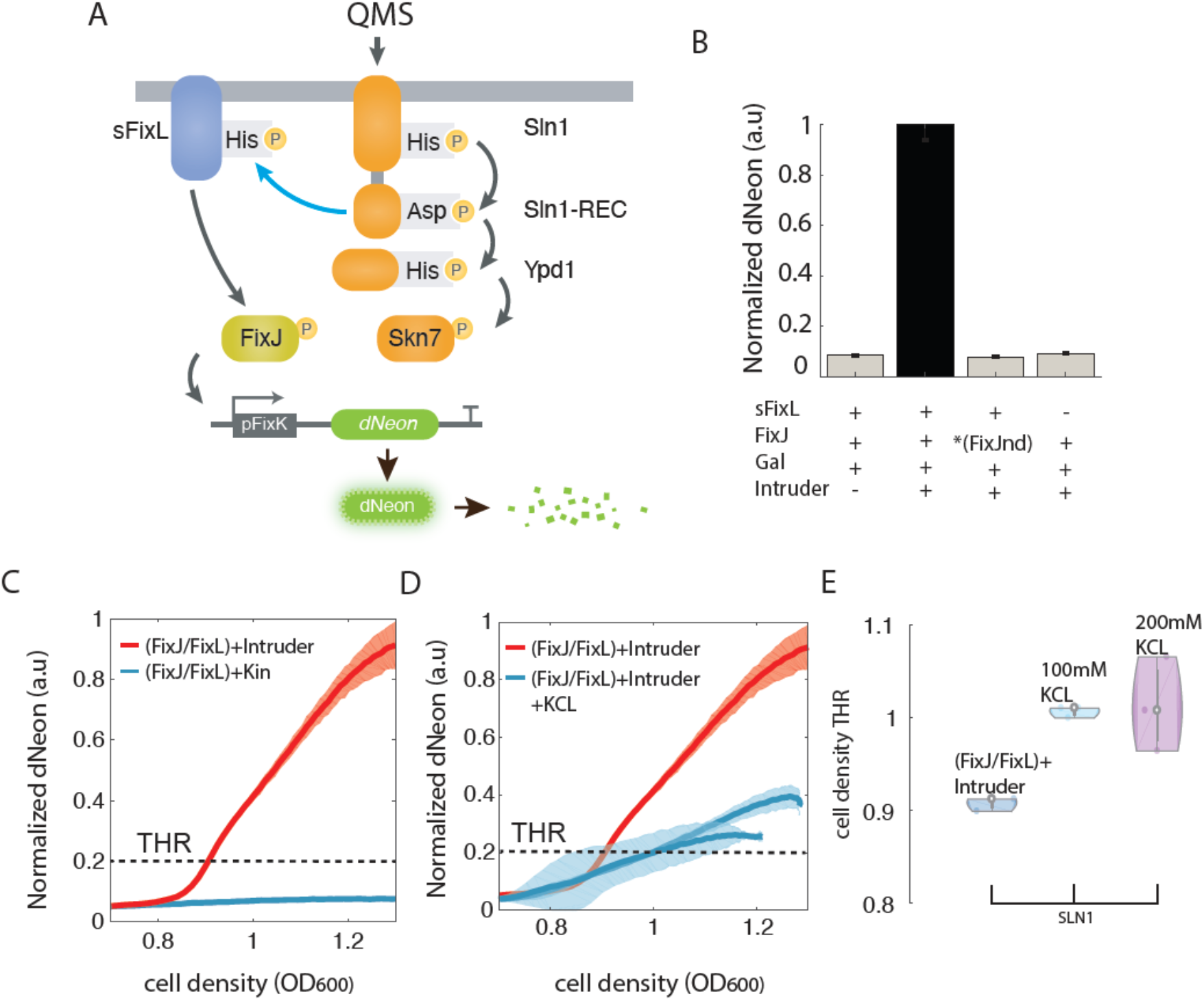
Plugging synthetic circuit to QMS for controlling orthologous gene expression. **A**, Schematics of synthetic system design integrating upstream QMS mechanism (orange). Bacterial HK FixL(in blue) fused with the HAMP and transmembrane domains of yeast SLN1. FixJ receiver (in green) was fused to herpes-virus VP16 activation domain. The synthetic promoter containing FixJ operators (pFixK) controls dNeon reporter. **B**, dNeon Reporter activation level in different configurations of synthetic circuit. FixJnd stands for non-phosphorylatable FixJ_nd_ mutant (D54Q) orthologous to SKN7(D427N). **C,** dNeon kinetics in cocultures of Kin with synthetic Fixj/FIxL system with Kin(blue curve) or Intruder (red curve). **D**, Hyperosmotic treatments resulted in the inhibition of FixL/FixJ system response as in the native QMS mechanism. **E**, THR shift to higher values caused by hyperosmotic stress. QMS curves show mean trends with 95% confidence intervals collected from three independent replicates.

We tested if QMS-activated SLN1 would directly transfer active phosphoryl-group to chimeric FixL which would then activate the FixJ transcription factor (Fig. 4A). Firstly, we checked whether the presence of FixJ and reporter alone would result in leaky reporter expression. Our quantifications suggest that FixJ alone is not phosphorylated by the native YPD1 phosphoryl group transfer mechanism (Fig. 4B, Fig. S10C), confirming the suitability of this bacterial HK system for orthologous pathway connecting QMS to any gene of interest. Indeed, using the full system (‘FixJ/FixL’) that includes FixL chimera showed clear activation that was dependent on QMS and FixL (Fig. 4B). To mimic orthologous SKN7(D427N) unphosphorylated mutant^21^ we mutated phosphor-accepting aspartate residue in FixJ (Fixj_*nd*_) completely blocked the reporter response (Figs. 4B and S10B, C). Because QMS does not seem associated with cell wall integrity and hypoosmotic responses (Fig. 1D), we tested whether imposing cell wall stress through applications of chitin-binding dye calcofluor white would trigger reporter response in ‘Kin’ without the presence of ‘Intruder’ in a synthetic signaling route. In this scenario, the reporter remained silent, indicating that cell wall stress may not trigger a QMS response (Fig. S10D). Finally, we investigated if the FixJ/FIxL strain responds to SLN1-deactivation by hyperosmotic conditions. Indeed, we found that KCL inhibited FixJ reporter response (Fig. 4D, E) in a similar manner to that observed in a native system (see Fig. 2C, D), that is by delaying a response, increasing THR and reducing response amplitude (Fig. 4E).

Altogether, these findings indicate that QMS can act through the orthogonal intracellular signaling pathway to control gene expression dynamics in the response to mechanical contact-based and cell density-dependent cues.

## Discussion

The understanding of molecular mechanisms underlying social interaction through QS in microbes is a key to controlling pathogenic infections and controlling biofilm formation unlocking various synthetic biology applications. In the fungal kingdom, putative QS mechanisms remained largely unknown^4,5^. We revealed a previously unidentified QS-like behavior that involves kin detection and mechanical contacts in the cell wall of baker’s yeast as opposed to classical chemical perception^1^. We showed that this QMS molecular mechanism depends on adhesin FLO11 activity which in turn inhibits SLN1, thus adjusting intracellular signaling for downstream gene expression in the presence of non-kin intruder strain. Heterologous contacts resulting from the difference in FLO11 expression competence between strains trigger QMS to resolve growth competition producing complex growth patterns reminiscent of flower petals. Furthermore, this QMS mechanism does not seem to involve hyperosmotic nor cell wall stresses through any of the canonical pathways, revealing the presence of an unknown signaling route mediated by SLN1. Also, these results are in strong agreement with a previous study that suggests an involvement of cell wall mannoproteins in the regulation of SLN1 activity^21^.

We demonstrated that the extracellular domain (ECD) of FLO11 tunes the response sensitivity and QMS density detection threshold. There is an intriguing possibility that highly glycosylated FLO11 ECD which can extend to the adjacent cells, could interact with N-glycans of SLN1 ECD, thus controlling SLN1 inhibition. An independent study should address the structural nature of FLO11-SLN1 interaction. There are mechanistic similarities between the proposed QMS and cis-dependent inhibition of delta-notch receptors in animal cells^51^, indicating possible evolutionarily convergence between fungi and animal cells in Kin cell detection strategies. Finally, we showed that the QMS mechanism can be used as a modular regulatory pathway to control gene expression through an orthogonal signaling cascade.

Our study opens opportunities for the development of new Kin detection systems that can be used to tailor heterologous biofilm dynamics for various applications including biomaterial and preventing biofilm-mediated pathogen infections. It remains a fascinating question whether a similar QMS-like mechanism could also operate in plants that use analogous HK osmosensing signaling circuits^14,17,52^ as such mechanisms would act in specific kin or cell-type detection and resolve mechanical growth conflicts during morphogenesis.

## Author contributions

- Conceptualization: KW, MGN
- Methodology: MGN, DRS. MA, ASM, SGR
- Investigation: MGN, DRS. MA, ASM, SGR
- Visualization: MGN, DRS
- Funding acquisition: KW
- Project administration: KW
- Supervision: KW
- Writing – original draft: KW, MGN, DRS
- Writing – review & editing: KW, MGN, DRS

## Data and materials availability

All data are available in the main text or the Supplementary Information.

## Materials and Methods

### Plasmids and gene constructions

Plasmids were constructed using isothermal Gibson assembly cloning^53^. A middle-copy (~10-30 copies) episomal vector pGADT7 (Takara Bio Inc.) was used as the basic expression backbone for the introduction of chimeric proteins, synthetic genes and reporter constructs. The DNA fragments used for the construction of sFLO11, synZIP, and sFixL/FixJ/ pFixK and dNeon reporter plasmids (including backbone and inserts) were either amplified from yeast genomic DNA using a polymerase chain reaction (PCR) or synthesized through Integrated DNA Technologies (IDT) services. Specific adapters at the 5’ and 3’ ends of DNA fragments were designed to facilitate efficient Gibson assembly and minimize secondary structure formation. High-fidelity Phusion PCR Master Mix with HF Buffer (Thermo Scientific, USA) was used for all PCR reactions, following the manufacturer’s protocol. The size of the amplified PCR products was confirmed in 1% agarose gel electrophoresis, purified using the EasyPure PCR Purification Kit (TransGen Biotech, China), and concentrated to approximately 100 ng/μL before Gibson assembly. Plasmids and DNA parts are summarized in Tables S1 and S2.

### Yeast genomic integrations and Flo11 knockouts

To study ‘Intruder’ presence in microfluidic and agar plate experiments, the constitutively expressed mCherry reporter was integrated into the genome of BY4741 and selected with G418 antibiotic (250mg/L). The correct insertion was confirmed by sequencing and fluorescence microscopy. To generate Δflo11 CEN.PK2-1C strain (‘Kin’ Δflo11), gRNA was designed to target the 5’ of FLO11 exon (sequence:” CGGATTTCCCAGGCTTCTAT”) by minimizing off-side effects and cloned into gRNA shuttle plasmid provided by Easy-Clone Marker free kit^54^ https://www.addgene.org/kits/borodina-easyclone-markerfree/). Then, the ‘Kin Δflo11’ mutant was obtained with CRISPR-based genomic knockout^54^ and confirmed by sequencing. The Galactose-inducible full-length FLO11 ORF clone with URA selectable marker was a kind gift from Dr. Jean Marie Francois (Institut National des Sciences Appliquées de Toulouse: Toulouse, FR). All episomal and integration plasmids were transformed in ultra-competent cells from E. coli DH5a strain using standard protocols.

### Yeast growth, transformation and culturing conditions

Yeast strains were cultivated in Yeast Nitrogen Base (Formedium, UK) plates supplemented with 2% of glucose at 30 °C. They were then grown in liquid low fluorescence Yeast Nitrogen Base media (Formedium, UK) supplemented with 2% sucrose at 30 °C in a shaking incubator. Minimal drop-out media compositions were used at all stages to prevent plasmid loss. Appropriate antibiotics were added to key dominant selection markers (e.g., G418 (250mg/L) or Hygromacin B (300mg/L)). The activation of the system was monitored using mNeonGreen fluorescent protein was fused to non-cleavable ubiquitin UbG76V for fast degradation^55^ referred to as the dNeon reporter. Plasmids were transformed into competent S. cerevisiae BY4741 and CEN.PK2-1C cells (a kind gift from Dr. Luis Rubio). Competent yeast cells were prepared and transformed with the Frozen-EZ Yeast Transformation II Kit (Zymo Research, USA), following the manufacturer’s instructions. All strains were transformed with three plasmids, each containing a selection marker for the complementation of either uracil, histidine, or leucine auxotroph. Transformant cells were selected on Yeast Nitrogen Base (Formedium, UK) 2% agar plates deficient in the corresponding selection nutrients. Engineered yeast strains are presented in Table S3.

### Microfluidics device fabrication and cell loading and imaging protocols

Fluorescence microscopy images of the engineered strains were acquired in 40 h time-lapse experiments using microfluidics chips to grow colonies of cells under constant perfusion of medium according to protocols previously described^56^. Cocultures were grown for first 15h before imaging and then imaged every 10 min up to 40h. Each cell trap (500×500×8 microns) contains > 10000 cells. A general scheme of the microfluidics setup is provided in Fig. S5. The internal features of the chips (media channels and cell traps) were laser-printed on polystyrene sheets. Next, the sheets were heated to approx. 160 °C. The printed pattern shrunk and increased in height by approx. 10-fold, rendering a high-relief mold that was used as a template for soft lithography. After a thorough cleaning, the device mold was covered in PDMS silicone (Sylgard), and degassed in a vacuum pump for approx. 20 minutes, and cured overnight at 80 °C. The resulting PDMS wafers were cut to the final shape of individual chips and input ports were opened using a 0.7 mm biopsy puncher (World Precision Instruments). Cover glasses were cleaned in a sonic bath, thoroughly rinsed with water and ethanol, and blow-dried using a nitrogen gun. In order to bond the PDMS chips to the cover glasses, the surface of both the chip and the glass was exposed to oxygen plasma with a Corona SB plasma treater (ElectroTechnics) for about 15 seconds. Then, both surfaces were brought together and kept at 80 °C overnight. The microfluidics chip was vacuumed for at least 20 minutes to facilitate the loading process. Flexible transfer tubing lines (ColePalmer) with an internal diameter of 0.5 mm were cleaned with 70% ethanol and connected in sterile conditions to syringes containing fresh media and to output containers. The input syringes were placed in a height control device to regulate the flow by adjusting their relative height to the output containers. The lines were then plugged into the chip one by one, ensuring that the liquid reached all features of the chip and no air bubbles were trapped. A separate input line had been connected to a falcon tube containing a liquid culture of the desired strain at an OD_600_ of 0.2. The tube containing the cells was lifted until the cells began to flow into the chip at a significant rate and started seeding the trapping region. Once 10-20 cells were retained in each trap, the flow from the loading port was reversed by decreasing the height of the syringe to prevent the entry of bubbles during the experiment. During the experiment we kept low flow conditions to supply nutrients but do not impose stress on cells.

### Fluorescent reporter data acquisition, processing and analysis in agar-based and liquid microwell plate experiments

In coculturing experiments either agar or liquid cultures, overnight cultures were diluted to a total starting OD_600_ of 0.2 (for co-cultures, OD_600_ = 0.1 of each strain, unless different ratios were used see Fig. S7). In microwell plate experiments, additional compounds (mannose, KCl, Concanavalin A) at indicated concentrations were added. 180 μL aliquots were pipetted from these diluted cultures to a multi-well plates, and 20 μL of galactose stock dilutions were added to each well to reach a concentration of 0.25%. In cocultures of ‘Kin’ with media extracts from ‘Intruder’. ‘Intruder’ was grown over night at standard conditions (200ul) to the stationary phase and next day cells were separated by filtering (0.45-micron filtration) and 20ul were used in cocultures with ‘Kin’ as described above. The plates were incubated at 30 °C in a FLUOstar® Omega Plate Reader (BMG LABTECH) performing time-lapse measurements of OD_600_ and dNeon fluorescence intensity (λ_Ex_ = 488 nm; λ_Em_ = 515 nm) every 10 minutes for up to 24 hours. All data points from fluorescence plate reader were processed to remove media background signal and normalized to the maximum observed fluorescence level based on positive control conditions, yielding values in the range of 0-1 for more comprehensive comparison between experiments. All plots contain complete data points from three biological replicates per condition. All experiments were repeated on two separated days (technical replicates) in triplicates yielding similar results. Cell density thresholds (THR) was set at OD_600_ corresponding to the 20% of maximal dNeon reporter signal measured in all compared co-cultures. In agar-based experiments cocultures (1:1 ratio) were grown on 1% agar with standard minimal media as described in above for 5-7 days to allow for colony outgrowth competition and the formation of ‘Kin’ protrusions at the edges. Images were taken using GFP (λEx = 488 nm; λEm = 515 nm) and mCherry (λEx = 583 nm; λEm = 610 nm) channels. Pictures were taken with an automated inverted Leica DMi8 fluorescence microscope equipped with a Hamamatsu Orca Flash V3 camera. Images were analyzed in Fiji by quantifying fluorescence signal along the arcs fitted to colony edge as shown in Fig. 1E, taking as reference tips of ‘Intruder’ fluorescence signal (mCherry).

### Microscopy image acquisition during microfluidic experiments

Experiments were run over 40 h under a constant perfusion of fresh media inside the microfluidic chip. Each trap was imaged every 10 minutes using three different channels: differential interference contrast (DIC), GFP (λEx = 488 nm; λEm = 515 nm) and mCherry (λEx = 583 nm; λEm = 610 nm). Pictures were taken with an automated inverted Leica DMi8 fluorescence microscope equipped with a Hamamatsu Orca Flash V3 camera. The image acquisition was controlled by the software μManager. Images were captured using a 20x dry objective with NA=0.8 (Leica, Germany). Fluorescence microscopy images were generated using a CoolLed pE600 LED source and a standard epifluorescent filter set (Chroma, USA). The multichannel image sequences were afterwards processed and analyzed using the Fiji software^57^, and custom R-studio and phyton scripts as described previously^56^. Briefly, each image was divided into 25 regions of interest (ROIs) and analyzed separately to isolate regions where cells were actively growing and could be tracked over time. The posterior analysis was done with custom R-studio scripts. Firstly, raw data were detrended using the detrend function from “pracma” R-studio v4.0.3 package and then smoothed with Savitzky-Golay Smoothing function (savgol), from the same package, with a filter length of 15 was applied and the signal was normalized between 0 and 1. To plot growth competition curves we quantified mCherry signal (‘Intruder)’ across each co-culture region (traps) in two different time points 15h after the start of experiment and a day later when experiment was completed. Generally, 24h was sufficient time for ‘Kin’ to outcompete ‘Intruder’ and decrease ‘Kin’ presence by nearly 50%.

