## Supplementary Information for "A quorum of mechano-sensing fungal consortia"

**Contains:**

- Figs. S1 to S10
- Tables S1-S3
- Video S1 with caption

**Figures S1-S10**

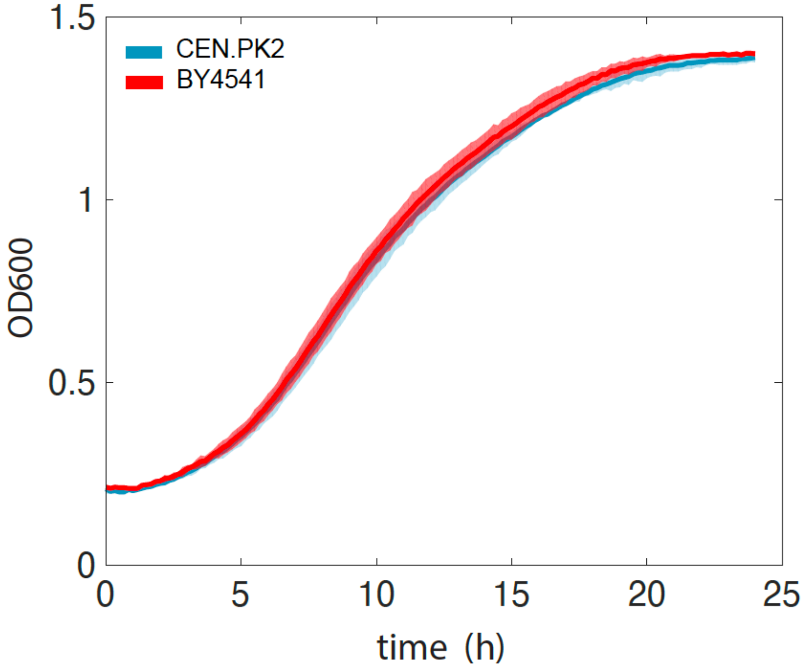


**Fig. S1. Comparison of growth kinetics in BY4741 and CEN.PK2 strains**

Both CEN.PK2-1C(blue curve) and BY4741 (red curve) show indistinguishable $OD_{600}$ growth kinetics under growth conditions used in the study. Curves show mean trends with 95% confidence intervals collected from three independent replicates.

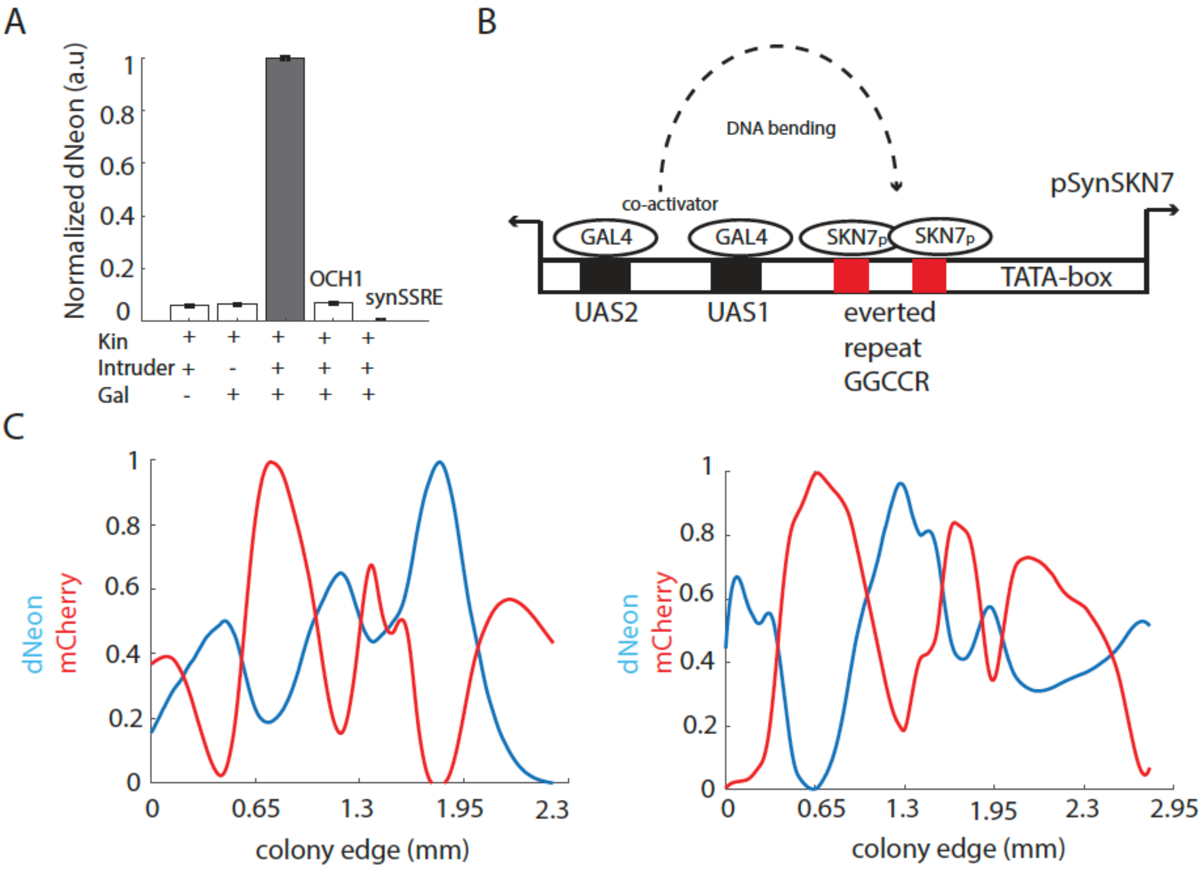


**Fig. S2. Design and testing of synthetic SKN7 responsive promoter and spatial patterns of growth in yeast consortia.**
**A**, Normalized dNeon activation with pSynSKN7 promoter and OCH1-based promoters. Note that the observed dNeon response is not triggered in configuration with pOCH1 or synthetic OCH1 variant promoters (psynSSRE). **B**, Schematic of synthetic promoter (pSynSKN7) design used to track ‘Kin’ activation dynamics. Two everted repeats of the SKN7-binding motif were placed upstream of the minimal promoter (TATA-box). Two GAL4-UAS sites were placed upstream with reverse orientation relative to the minimal promoter. The proximity of UAS sites (black boxes) could facilitate SKN7-dependent activation of pSynSKN7 promoter possibly by co-activation or DNA bending. However, galactose addition alone does not activate the promoter, indicating a requirement for the proximity of active SKN7 bound to GCCR sites (red boxes). **C**, Other examples of mutually excluding binary patterns of ‘Kin’ and ‘Intruder’ growth regions on the edge of the mixed colony. Plots represent dNeon (blue) and mCherry(red) average signals across the edge of the colony (arc) as indicated in Fig. 1E.

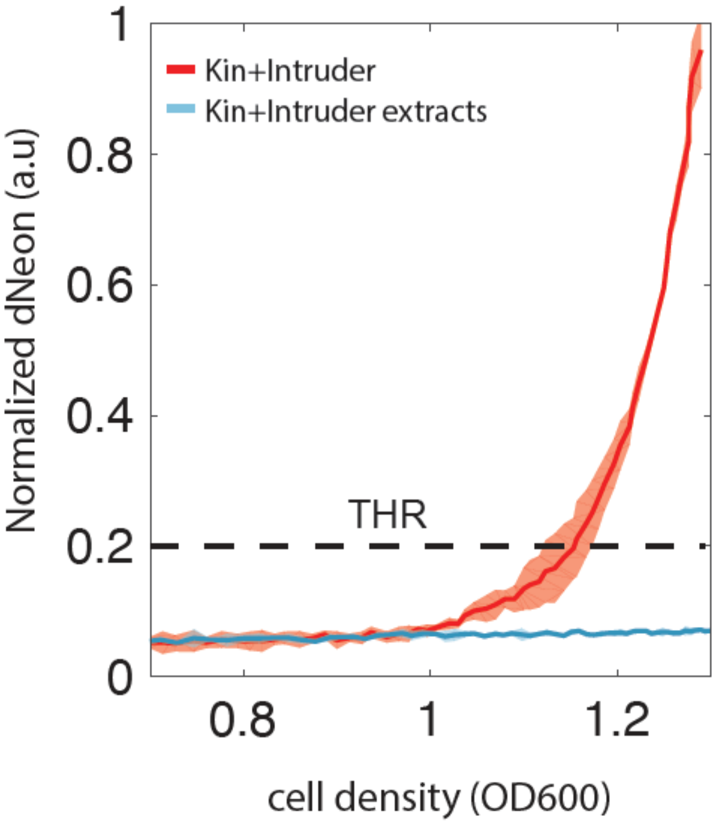


**Fig. S3. dNeon reporter activation is not mediated by chemical cues originating from 'Intruder'**

'Kin' was cocultured with media extracts in which 'Intruder' was previously grown to the stationary phase. These extracts could potentially contain QS-like molecules released by 'Intruder' cells. However, dNeon reporter showed no activity under this condition (blue curve) compared to control 'Kin'+'Intruder' (red curve). This data indicated that the signal is likely not of chemical nature. QMS curves show mean trends with 95% confidence intervals collected from three independent replicates.

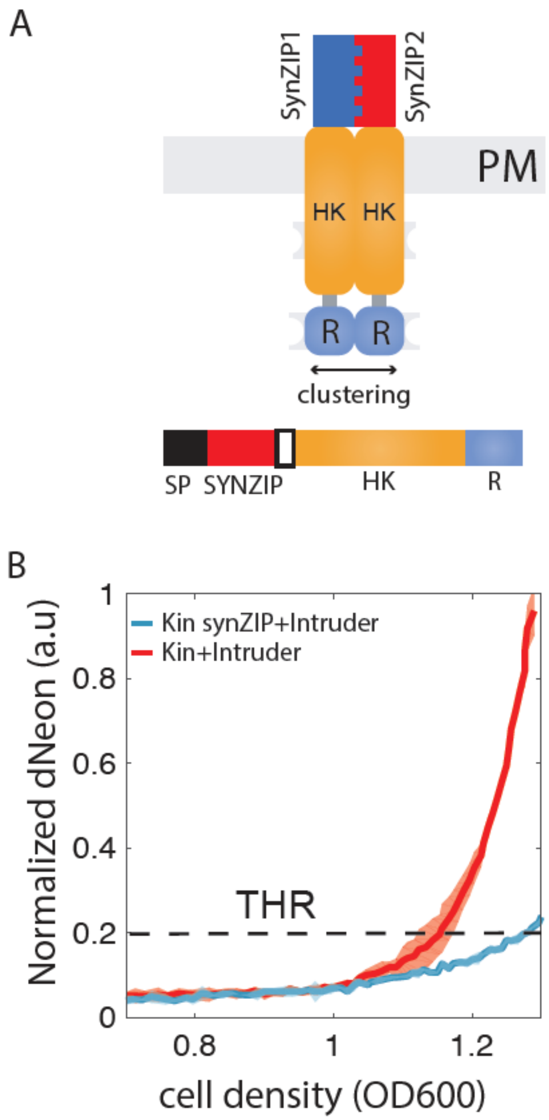


**Fig. S4. Deactivation of SLN1-pathway by forcing extracellular SLN1 dimerization with synthetic zippers**

**A,** Schematic of engineered SLN1 designs. Signal peptide (SP) has been fused to SynZIP1 or SynZIP2 zippers[35] following Transmembrane and HK, and Receiver (R) domain of SLN1. This configuration allows for a direct dimerization of SLN1 through extracellular domains. **B**, QMS curves in controls and engineered 'Kin synZIP' coculture with 'Intruder'. Note that imposing extracellular dimerization of SLN1 attenuated dNeon response by ~5-fold. QMS curves show mean trends with 95% confidence intervals collected from three independent replicates.

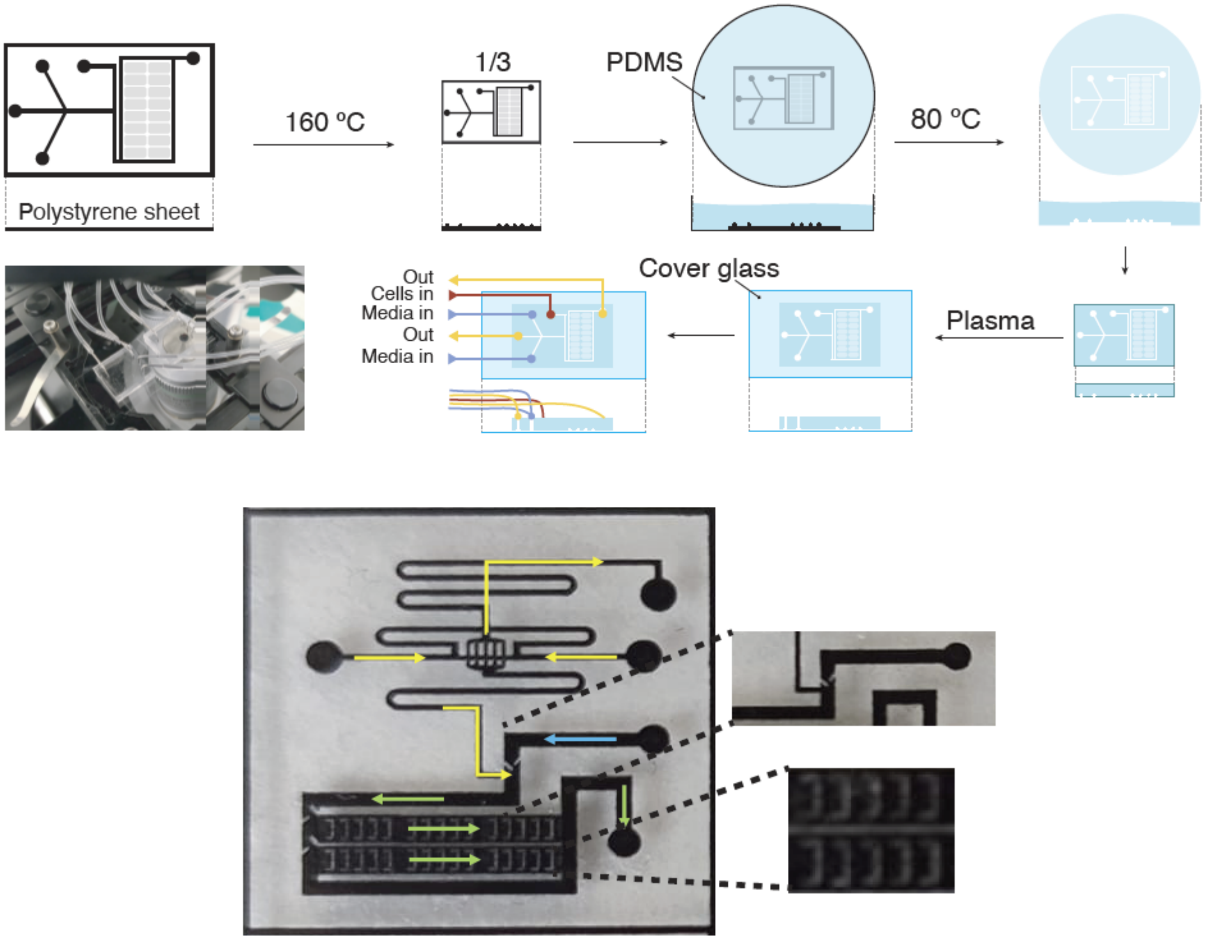


**Fig. S5. Microfluidics setup.**

Upper panel show the process of microfluidic device fabrication[56]. The design of the chips is laser-printed on polystyrene sheets, thermally treated, and used as a mold for soft lithography with PDMS. The chips are cut and plasma-bonded to a cover glass. Ports are opened on the surface of the chip with a biopsy puncher, and input and output lines are plugged. Middle panel depicts inlets and outlets on the chip used during microfluidic experiments. Bottom panel, show typical experimental setup. Initially, cells are loaded through cell port (blue line) after they seeded the traps the flow is reversed (yellow arrows) and kept at low rate to avoid cell stress. Channel widths were 120µm for in the mixer module and 500µm in main channels. The approximate height of channels was 25 microns. Cell traps had 500µm x 500µm size and height of approximately 8 microns.

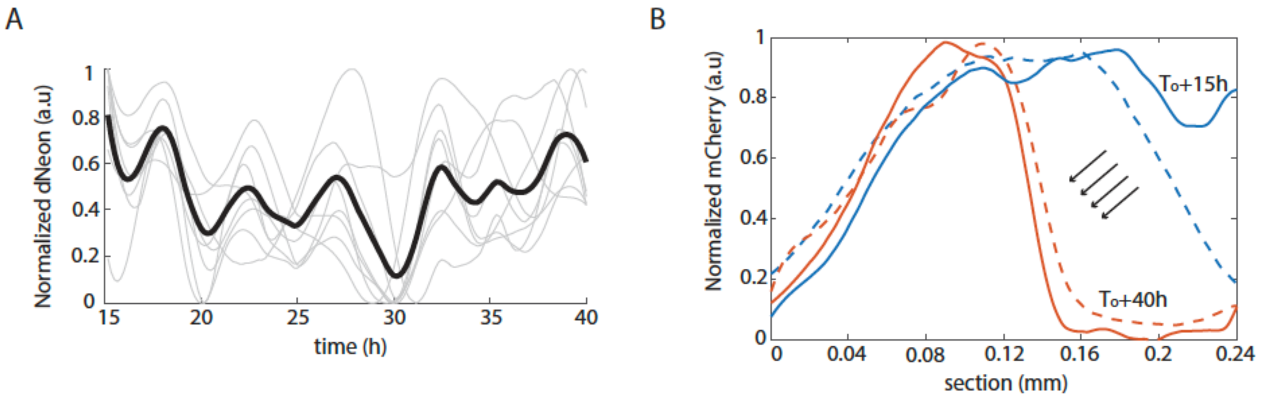


**Fig. S6. Spatial-temporal analysis of QMS dynamics.**

**A**, Time-course plots from microfluidic experiments of ‘Kin’ and ‘Intruder’ cocultures in 10 independent trapping regions (n=10, each trapping region contains > 10000 cells), and average trace is shown in black. Note the transient switch-like dynamics of ‘Kin’ in response to the presence of ‘Intruder’. **B**, Other examples of the time evolution of growth competition between ‘Kin’ and ‘Intruder’ in microfluidics traps. Only intruder presence is shown with mCherry reporter. Note that over the course of ~24h (red curves) ‘Intruder’ occupancy drastically decreases to nearly 50% less of its original size caused by outgrowing ‘Kin’ population. Color coding is as in Fig. 2G.

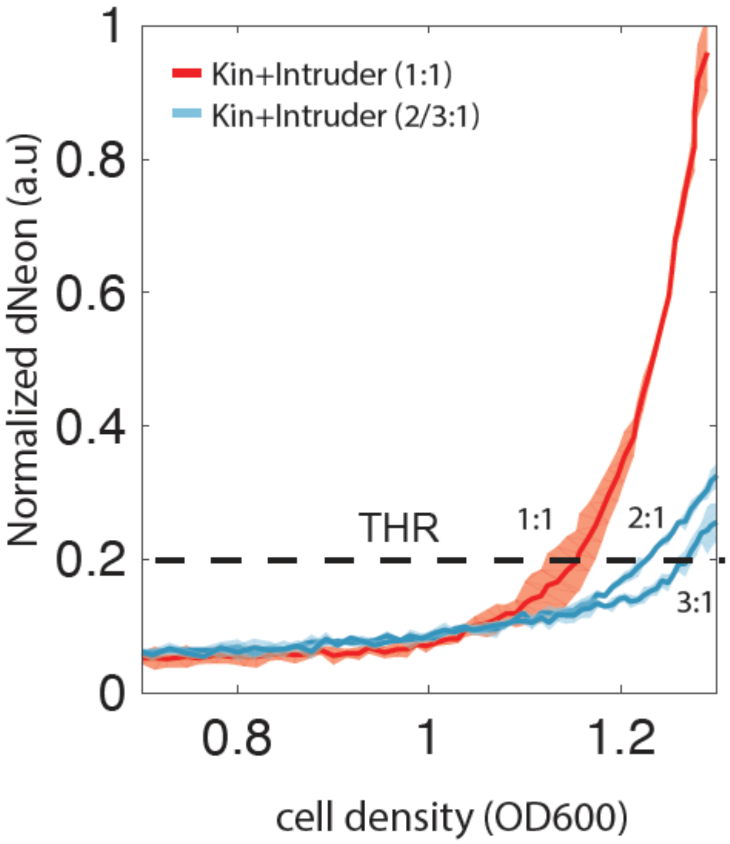


**Fig. S7. Effect of increasing the Kin: Intruder ratio in cocultures.**

‘Kin’ was inoculated with ‘Intruder’ in the following ratios (1:1, 2:1, and 3:1). Progressive increase of ‘Kin’ (and also FLO11) with respect to ‘Intruder’ shifted THR to higher values, indicative of stronger inhibition of SLN1 pathway. QMS curves show mean trends with 95% confidence intervals collected from three independent replicates.

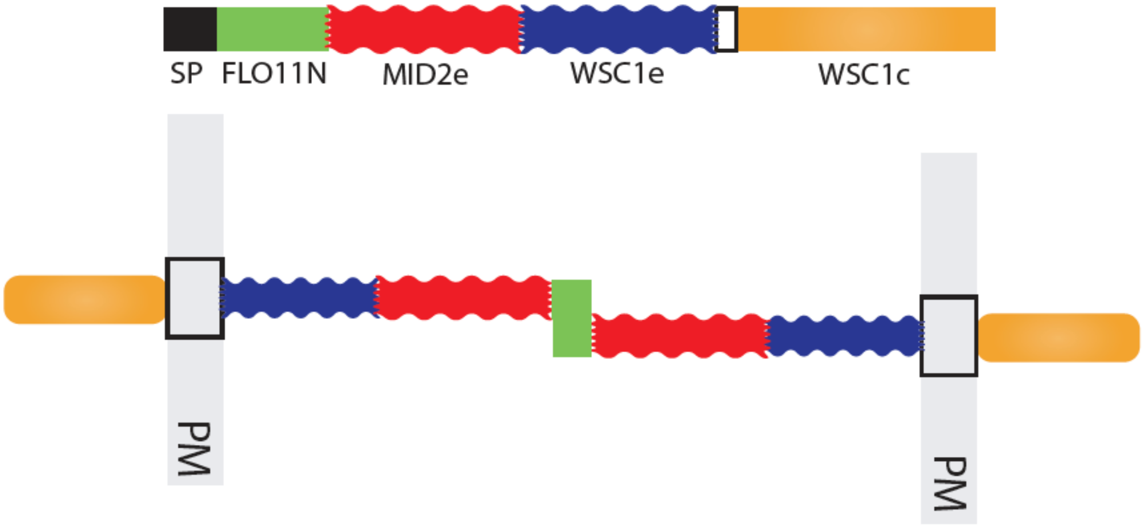


**Fig S8. sFLO11 construction schematic.**

Signal peptide and FLO11 extracellular region (1-211) were fused to extracellular domains of MID2 (43-219) and WSC1 (22-245) following transmembrane and intracellular domains of WSC1 (245-368). This chimeric construct extends to the cell wall of neighboring cells > 100nm, providing the possibility for either cis and trans interactions between sFLO11 in adjacent cells.

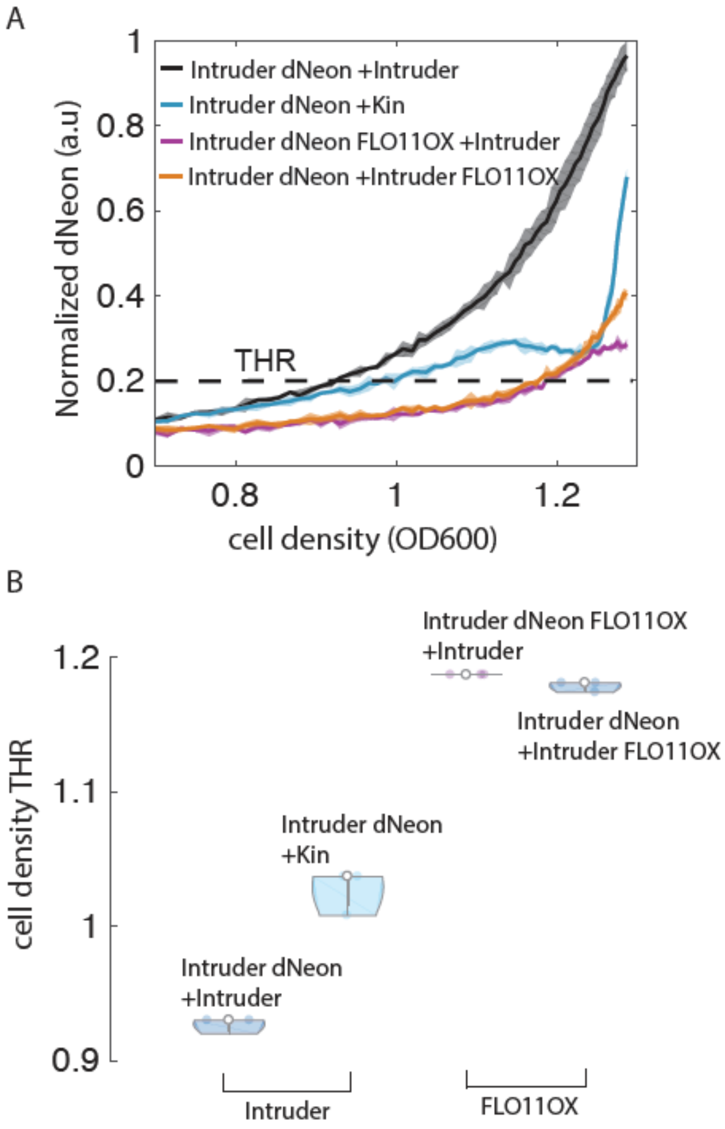


**Fig S9. Inhibition of Intruder's QMS response in cocultures with Kin and FLO11 overexpressing 'Intruder'.**

**A,** QMS kinetics in 'Intruder' is triggered by the introduction of Kin or FLO11 in 'Intruder'. Note a progressive repression of dNeon reporter signals in the presence of FLO11. QMS curves show mean trends with 95% confidence intervals collected from three independent replicates. **B**, cell density THR changes are dependent on the presence of FLO11 adhesin.

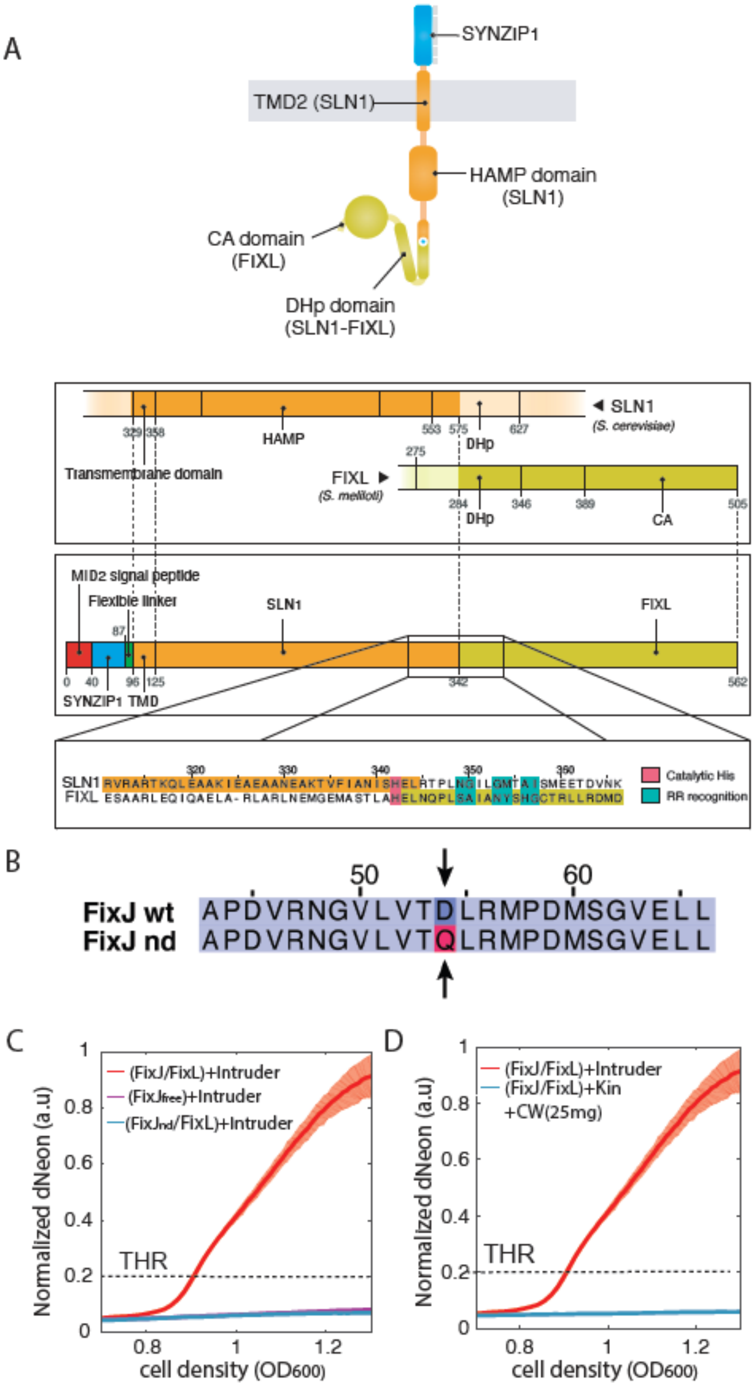


**Fig. S10. Synthetic QMS-driven system in yeast based on FixL/FixJ bacterial HK system.**

**A**, MID2sp(signal peptide)-SynZIP1 was fused to the transmembrane domain (TM) of SLN1 using a flexible GS linker followed by SLN1 HAMP domain connected through the complementary helical region to HK part of FixL composed of DHp and CA domains[46] based on secondary structure alignment. In this configuration, sFixL receives the phosphoryl group from the receiver domain of native SLN1 (see Fig. 4A, B). FixJ transcription factor was fused to tandem repeats of VP16 (VP64) transactivation domain of herpes simplex virus[48]. **B**, Sequence annotation of FixJ dephosphorylating/deactivating mutation D54Q orthologous to SKN7(D427N). **C,** dNeon reporter time-lapse kinetics in $FixJ_{nd}$(D54Q) mutant. **D,** Activation of 'FixJ/FixL' strain does not depend on cell wall stress caused by calcofluor white(CW) treatments. QMS curves show mean trends with 95% confidence intervals collected from three independent replicates.

**Table S1. Summary of plasmids generated in this study**

| Construct name | Components | Auxotrophic markers (yeast) | Notes |
|---|---|---|---|
| dNeon reporter | pSynSKN7 :: UbG76V-mNeonGreen | HIS | Main reporter plasmid used in this study |
| OCH1 reporter | pOCH1 :: UbG76V-mNeonGreen | HIS | OCH1 gene promoter |
| synSSRE reporter | psynSSRE :: UbG76V-mNeonGreen | HIS | psynSSRE synthetic promoter |
| STL1 reporter | pSTL(1-1000) :: UbG76V-mNeonGreen | HIS | STL1 gene promoter fragment responsive to osmotic stress |
| RLM1 reporter | pRLM1(1-864) :: UbG76V-mNeonGreen | HIS | Promoter Fragment of RLM1 downstream of wall stress response pathway |
| ZIPsln (SynZIP1/2) | pGAL7 :: SP(SLN1)-SynZIP1/2-SLN1(TMD)-SLN1cyt | URA(SynZIP1), LEU(SynZIP2) | This study Chimeric SLN1 protein where ectodomain has been replaced with complementary synthetic leucine zippers |

| | | | |
|---|---|---|---|
| sFLO11 | pGAL1 :: FLO11N(1-211)-MID2ecto(43-219)-WSC1ecto(22-245)-WSC1TMD-WSC1cyt(245-368) | URA | This study Signal peptide and FLO11 extracellular region (1-211) were fused to extracellular domains of MID2 (43-219) and WSC1 (22-245) following transmembrane and intracellular domains of WSC1 (245-368). |
| pFixK | Minimal TATA + FixJ binding sites[48] | HIS | This study Synthetic yeast promoter with FixJ operator sites |
| sFixL | pGAL7 :: SP(MID2)-SynZIP1-SLN1TMD-SLN1HAMP-linker- FixLHK | URA | This study<br><br>MID2sp(signal peptide)-SynZIP1 was fused to transmembrane domain (TM) of SLN1 using flexible GS linker followed by SLN1 HAMP domain connected through the complementary helical region to HK part of FixL composed of DHp and CA domains based on secondary structure alignment. |
| FixJ<br><br>$FixJ_{nd}$ mutant | pGAL7 :: VP64-FixJ<br><br>pGAL7 :: VP64-$FixJ_{nd}$ | LEU | This study FixJ fused to VP16 repeats (VP64) |

| | | | |
|---|---|---|---|
| | | | transactivation domain |
| eLeu | Empty insert pGADT7 | LEU | This study |
| eUra | Empty insert pGADT7 | URA | This study |
| eHis | Empty insert pGADT7 | HIS | This study |

### Table S2. Summary of DNA sequences

| DNA fragment | Sequence |
|---|---|
| pSynSKN7 | tcatgttgatgctctgcataataatgcccataaatatttccgacctgcttttatatctttgctagccaaactaactgaacatagctacacattattttcagcttggctattttgtgaacactgtatagccagtccttcggatcacggtcaacagttgtccgagcgctttttggaccctttcccttatttttgggttaaggaaaatgacagaaaatatatctaatgagccttcgctcaacagtgctccgaagtatagctttccaaaaggaactggccgtcgttttacagccttcatgctccttgatttccctagctcagtcctaggtatagtgcggaaatcaaggagcatgaaggctgtaaaacgacggccagtatcgcagcccctgtccctatcagtgatagagaacgtataaggagtttactccctatcagtgatagagatattacttcttattcaaatgtaaggcatatatatatgtgcgcgtatatacatgattatatggcatgtatgtgctctgtatgtatataaaactctttttttctttttttctctaaattttttttccttatacattaggacctttgcagcataaattactatacttctatagacacgcaaatacaaatacacacactaa |
| UbG76V-mNeonGreen | atggtgagcaaaggagaagaagataatatggcgagcttaccggccactcatgagctgcatattttcgggagcatcaatggagtcgattttgacatggtcgggcagggcactggaaacccgaatgacggctatgaagagctaaatctgaagtcaacgaagggtgatttgcagttctccccgtggatcttggtgcctcacataggatatgggtttcatcagtatttgccttatcccgacggtatgagtccctttcaagcggctatggtagacgggtctgggtaccaggttcacaggacgatgcaatttgaagacggggcttctcttactgtgaactatcgttatacttatgaagggagccacatcaaaggcgaagctcaggtcaagggaactggcttccccgcggacggccctgtaatgacgaactccttgacggcggccgattggtgcagaagcaagaagacatatccaaatgataagacgatcatttcaacttttaagtggagctacacaactggcaatgggaagagatatcgttccacagcccgtactacatatacctttgcaaagcctatggcagcgaactatctaaaaaatcagccgatgtatgtatttagaaaaacagagctaaagcatagtaagacagaactgaatttcaaggagtggcaaaaagcatttaccgacgtgatgggtatggatgaactatataaa |
| pOCH1 | ctttacgcctctctttattctttttgggtaaattgcttaaactatttggccggcccaccgcgaaaagatttggctgggcctcaactaaacgcgcctttttggacttttcacgttgcagggacagcaacgtcaaaacttctgcattaaggtagtttggtagcttggtagccactttagtatttctgccttcttcgaataccgacattatttctcgccaatccacattctctctccccatctgcatccttttatatttaatagggataggttgttttagttctttgattccgttttcatttcaagagcaataatagcaatttggaaaaagaaagcaagtaaaagaaagaagagatc |
| pRLM1 | gatatcttaagacctgagttaccgagtcttacctgcttaactaaaaccatttctgccgcctgatgtgggtcttccgtaacagccaaattacatagttggcaagacaattaatatatgatacctctgttctctgggtcctaatatatatatatatatatatatatatacgaaaaggctgagaagaataaggcagatgttagcttagcttcttttactcaagcaatgaaaacaaaatcgaagggaaacaaattcaaaccaagcggccgagttatctcccgttcctaacctactacactaatcacatttttccgtggaaataacgttattgaaaatgcgttcttgcctgccatacgtgggtgaaatacttccacgttcgcaaaaaatacttccacggtgacgaagtctgtctcagtcgtatattaaatgcagaaatcgtcttatcattattgggctctcttaacggcgcagcatcaccgggtgatgaatgccaagccgcagaaagaaagaaaaaaatttacttcagatttctgataaaaataaaacggaagagatgaaagctaataatagaaacagctcgatcttcctctgaacaataataattaaaggacagacaaaaagaaacgtaagaaagaagcgagcctgttctaaagtgttcaacgactgattcaattagaactgcctactcctgatagccaactcaacttttgactcgttaaagtaattgaaagctggcaagcagaattattcttttttttttcaaggtttctatcacgttgtgaggttaatatc |

| | |
|---|---|
| | ccccggagcaaacaggctgaagcgtgaaaaaaacttaaatattaaagtgtcgcaaa<br>actatactatagatacaac |
| pSTL1 | caatgattctgaaatactccttttacaacctttgcaaagataatgtctttcagtctgatattac<br>gagcgacctagaaccactaatccatattcttcattcaacttactccattttccttggcaaac<br>aatgccccacaatcatatacgtcataactataagggatatgtctggaatgcggccaag<br>atagaattaaagggctgcagaacaccactactgatactcattgccaaggctaggagg<br>caccatccgtttcattttctttgaggtaagccaatcatgaaatagtatacacatccataacg<br>gacgtacggacgaaataagtgccgttgtcccactattccaccgcatttggcccatttggc<br>tcactttgactcaacttgcgtcattttaactgatatgaagggtccgactttgtcctttttcggcc<br>accgcataccccacggcgatgcctccgctacctgcatttgagtagcatctccgtttcgcg<br>gggtattcggcgctacgtcgcctgttcgagcggctctgttcgttgcatgaaactaaaataa<br>gcggaaagtgtccagccatccactacgtcagaaagaaataatggttgtacactgtttctc<br>ggctatataccgtttttggttggttaatcctcgccaggtgcagctattgcgcttggctgcttcg<br>cgatagtagtaatctgagaaagtgcagatcccggtaagggaaacacttttggttcacctt<br>tgatagggctttcattggggcattcgtaacaaaaaggaagtagatagagaaattgaga<br>aagcttaagtgagatgttttagcttcaattttgtccccttcaacgctgcttggccttagaggg<br>tcagaattgcagttcaggagtagtcacactcatagtatataaacaagccctttattgatttt<br>gaataattattttgtatacgtgttctagcatacaagttagaataaataaaaaatagaaaaa<br>tagaacatagaaagttttagacc |
| psynSSRE | ctcgaatttggccggcccacaccggtagatttggctgggcctcgggcatatatatatgtg<br>cgcgtatatacatgattatatggcatgtatgtgctctgtatgtatataaaactctttttttcttttttt<br>ctctaaattttttttccttatacattaggacctttgcagcataaattactatacttctata |
| ZIPsln<br>(SynZIP1) | atgttatcatttacaaccaaaaattcctttaggttacttctattgattctgagttgcatttccacc<br>attagggcgcaatttttttgttcaaagtagctcctctaactcaagcgcggtctcaaacctggt<br>tgcacagcttgaaaacgaagtcgcatccctggaaaacgagaatgaaacgctgaaaa<br>aaaagaacttgcacaagaaagacttaattgcctatttggaaaaggagattgctaaccta<br>agaaaaaaaatagaagaaggttctgattctacttctggcagcaagctagcaaaaatca<br>tcaccggcactgtcatcgctattggtgtctttgtcattttgttaacccttcctctagcacactgg<br>gcagtgcaaccaattgtacgtctacaaaaggcaactgaattaattacagaggggaga<br>ggccttcgaccgagcactccaagaacgataagcagagccagttcattcaaaagagg<br>atttagttctggatttgctgttccttcttcgttattacaatttaatactactgaagctggcagca<br>ccacaagcgtaagtggccatggaggcagtgctcatggcagtggtgcagctttttcagca<br>aatagtagcatgaaaagcgctataaaccttggaaatgagaaaatgtcacctccagag<br>gaggagaacaaaataccgaataaccataccgatgctaaaatatcaatggatggctcg<br>ctaaatcacgatttgcttgcaccacattccttgagacataatgacactgacagaagttcc<br>aatagatctcacattctcacaacttctgcaaatttaactgaagctaggctaccagattata<br>gaagactattttctgatgaactttccgatttaacagaaaccttcaatactatgacagacgc<br>attagaccaacattatgctcttttagaagaaagagttagggcgaggacaaaacaactc<br>gaagctgccaagattgaggcagaggctgcaaatgaagcaaaaaccgtctttattgcc<br>aatatttcgcatgaattgagaacacctttaaatggtattctgggcatgacggctatttcaat<br>ggaagaaaccgatgttaacaaaataagaaatagtttaaaactcatttttagatcaggtg<br>agcttttgcttcatattctaacggaattgttaactttttccaaaaacgttcttcaaagaacga<br>aactggagaaaagagatttttgcattaccgatgttgccttacaaataaagtcaatatttgg<br>taaagttgcaaaggatcagcgtgttcgtctttcaatatcattgtttcctaatttgataaggac<br>aatggttctttggggtgattccaacagaattattcagattgtgatgaatctagtgtccaatgc<br>actaaagttcacccctgtagatggtaccgttgatgtaagaatgaaactgttgggtgaata<br>cgacaaagaattaagcgagaagaagcaatacaaagaagtgtatatcaaaaagggt |

| | |
|---|---|
| | acagaagtaaccgaagatttagaaactacagataaatacgatcttccaactttatcgaa<br>ccataggaaaagtgttgatttagaatccagcgctacttccctaggaagtaatagagaca<br>cttcgacaattcaggaagagataacaaaaagaaatactgttgcgaatgaaagtatcta<br>taagaaagtgaatgagagggaaaaagcttcgaatgatgatgtatcttctatagtatcaa<br>caactaccagctcgtatgataacgctatcttcaatagtcagttcaataaagcacttggctc<br>agatgatgaagaaggtggtaacctaggaagacctatcgaaaatcccaaaacatgggt<br>tatttctattgaagtggaagacactgggcctggtattgacccttccttacaagaatctgtatt<br>tcatccatttgttcaaggtgatcaaacattgtccaggcaatatggtggtactggcttaggtc<br>tatcaatctgtagacagttagcaaatatgatgcatggaacgatgaaattagagtcgaaa<br>gtaggtgttggtagtaaattcacttttaccttgccattaaatcaaactaaagagatcagtttt<br>gcagatatggagtttccttttgaggacgaatttaatcctgagagtagaaagaatagaag<br>agtcaagtttagtgttgctaaaagcatcaagagccgacaatccacatcatctgttgcaac<br>accagctacaaatagaagtagcctaaccaacgacgtgctaccggaggtaagaagta<br>aaggtaagcatgagacgaaagatgttggaaatcctaacatgggaagagaagaaaa<br>aaacgacaatggagggcttgaacaactgcaggaaaaaaatattaaaccttctatatgt<br>cttacaggtgctgaagttaacgaaaaaaattccttgtcttctaagcatcgttctcgacatg<br>aaggtctaggttctgtcaatcttgatagaccatttttgcaaagtactggtacagccacatc<br>gagtagaaacatccccacagtcaaagacgacgataaaaatgaaacaagtgtcaaa<br>attttggttgtagaagataatcatgtaaatcaggaagttatcaaaagaatgttgaacttgg<br>agggcattgaaaatattgaactggcttgcgatggccaagaagcattcgacaaagttaa<br>agaattgacatctaagggcgaaaattataatatgattttcatggatgtccagatgcctaa<br>agtggatggtttactttctaccaagatgataaggcgcgatttaggttatacctcacccattg<br>tcgctctaaccgcttttgctgacgatagcaacattaaagaatgtttggaatcaggaatga<br>acggatttttatcgaaaccaatcaaaagaccaaaattgaaaactattcttactgagttttgt<br>gcagcatatcagggaaagaaaaataacaaatga |
| ZIPsln<br>(SynZIP2) | atgttatcatttacaaccaaaaattcctttaggttacttctattgattctgagttgcatttccacc<br>attagggcgcaatttttttgttcaaagtagctcctctaactcaagcgcggtctcagctagga<br>atgcgtaccttagaaagaagatcgccagattaaagaaagacaatcttcagctggaaa<br>gggacgagcagaacttggaaaaaattatagcgaatcttagggacgagattgctcgttt<br>agagaacgaagtcgcaagccacgagcaaggttctgattctacttctggcagcaagcta<br>gcaaaaatcatcaccggcactgtcatcgctattggtgtctttgtcattttgttaacccttcctc<br>tagcacactgggcagtgcaaccaattgtacgtctacaaaaggcaactgaattaattac<br>agaggggagaggccttcgaccgagcactccaagaacgataagcagagccagttcat<br>tcaaaagaggatttagttctggatttgctgttccttcttcgttattacaatttaatactactgaa<br>gctggcagcaccacaagcgtaagtggccatggaggcagtgctcatggcagtggtgca<br>gctttttcagcaaatagtagcatgaaaagcgctataaaccttggaaatgagaaaatgtc<br>acctccagaggaggagaacaaaataccgaataaccataccgatgctaaaatatcaa<br>tggatggctcgctaaatcacgatttgcttgcaccacattccttgagacataatgacactga<br>cagaagttccaatagatctcacattctcacaacttctgcaaatttaactgaagctaggcta<br>ccagattatagaagactattttctgatgaactttccgatttaacagaaaccttcaatactat<br>gacagacgcattagaccaacattatgctcttttagaagaaagagttagggcgaggaca<br>aaacaactcgaagctgccaagattgaggcagaggctgcaaatgaagcaaaaaccg<br>tctttattgccaatatttcgcatgaattgagaacacctttaaatggtattctgggcatgacgg<br>ctatttcaatggaagaaaccgatgttaacaaaataagaaatagtttaaaactcatttttag<br>atcaggtgagcttttgcttcatattctaacggaattgttaactttttccaaaaacgttcttcaa<br>agaacgaaactggagaaaagagatttttgcattaccgatgttgccttacaaataaagtc<br>aatatttggtaaagttgcaaaggatcagcgtgttcgtctttcaatatcattgtttcctaatttga |

| | |
|---|---|
| | taaggacaatggttctttggggtgattccaacagaattattcagattgtgatgaatctagtg<br>tccaatgcactaaagttcacccctgtagatggtaccgttgatgtaagaatgaaactgttg<br>ggtgaatacgacaaagaattaagcgagaagaagcaatacaaagaagtgtatatcaa<br>aaagggtacagaagtaaccgaagatttagaaactacagataaatacgatcttccaact<br>ttatcgaaccataggaaaagtgttgatttagaatccagcgctacttccctaggaagtaat<br>agagacacttcgacaattcaggaagagataacaaaaagaaatactgttgcgaatga<br>aagtatctataagaaagtgaatgagagggaaaaagcttcgaatgatgatgtatcttctat<br>agtatcaacaactaccagctcgtatgataacgctatcttcaatagtcagttcaataaagc<br>acttggctcagatgatgaagaaggtggtaacctaggaagacctatcgaaaatcccaa<br>aacatgggttatttctattgaagtggaagacactgggcctggtattgacccttccttacaa<br>gaatctgtatttcatccatttgttcaaggtgatcaaacattgtccaggcaatatggtggtact<br>ggcttaggtctatcaatctgtagacagttagcaaatatgatgcatggaacgatgaaatta<br>gagtcgaaagtaggtgttggtagtaaattcacttttaccttgccattaaatcaaactaaag<br>agatcagttttgcagatatggagtttccttttgaggacgaatttaatcctgagagtagaaa<br>gaatagaagagtcaagtttagtgttgctaaaagcatcaagagccgacaatccacatca<br>tctgttgcaacaccagctacaaatagaagtagcctaaccaacgacgtgctaccggag<br>gtaagaagtaaaggtaagcatgagacgaaagatgttggaaatcctaacatgggaag<br>agaagaaaaaaacgacaatggagggcttgaacaactgcaggaaaaaaatattaaa<br>ccttctatatgtcttacaggtgctgaagttaacgaaaaaaattccttgtcttctaagcatcgt<br>tctcgacatgaaggtctaggttctgtcaatcttgatagaccattttgcaaagtactggtac<br>agccacatcgagtagaaacatccccacagtcaaagacgacgataaaaatgaaaca<br>agtgtcaaaattttggttgtagaagataatcatgtaaatcaggaagttatcaaaagaatgt<br>tgaacttggagggcattgaaaatattgaactggcttgcgatggccaagaagcattcgac<br>aaagttaaagaattgacatctaagggcgaaaattataatatgattttcatggatgtccag<br>atgcctaaagtggatggtttactttctaccaagatgataaggcgcgatttaggttatacctc<br>acccattgtcgctctaaccgcttttgctgacgatagcaacattaaagaatgtttggaatca<br>ggaatgaacggatttttatcgaaaccaatcaaaagaccaaaattgaaaactattcttact<br>gagttttgtgcagcatatcagggaaagaaaaataacaaatga |
| sFLO11 | atgcaaagaccatttctactcgcttatttggtcctttcgcttctatttaactcggctttgggttttc<br>caactgcactagttccaagaggatcctccgaaggaactagctgtaattctatcgttaatg<br>gctgtcccaacttagacttcaattggcacatggaccagcaaaatatcatgcagtatacttt<br>ggatgtgacttccgtttcttgggttcaagacaacacataccaaatcactattcatgtcaaa<br>ggtaaagaaaatattgacctgaagtatctatggtctttgaaaatcattggtgtcactggtcc<br>aaaaggtaccgtccaactatacggttacaacgaaaatacctatttgattgacaacccaa<br>ctgatttcacagccacttttgaagtttatgccacacaagatgtcaacagctgtcaggtgtg<br>gatgcctaacttccaaattcaattcgagtatttgcaaggtagtgccgctcaatatgcaagc<br>tcctggcaatggggaactacatcttttgatttgtctactggttgtaacaactatgacaatca<br>aggccactctcaaacggatttcccaggcttctattggaacatagattgtgacaataattgt<br>ggcggtacgaagtcatctcgttcttccgtaagtagagttagttcttcaagttccattttgtcat<br>ccagtatggtttcttcctcaagtgctgactcatcttcccttacttcatcgacatcaagtaggtc<br>cctcgtgtcacatacgagttcgtctaccagcattgcctccatatcgttcacatcattcagttt<br>ctcatcagattcaagcaccagcagctcttcctctgcttcttcagattcttcatcctcctcatcc<br>ttttccatctcttcgacatccgcaacttctgaatcatcgacgtcttctacgcaaacgtcaac<br>atcatcttcatcctcactgtcgtcaacgccgtcctcttcatcatccccatcaacaatcacttc<br>tgcaccttcaacctcctccacaccatccactactgcctataatcaaggaagcactatcac<br>cagtattattaacggtaagacgattctctcaaatcattatactaccgttacgtacacaccat<br>cagcaaccgctgattcaagcaataaatccaaaagttcgggtctttcttatgaatacgtga |

| | |
|---|---|
| | attgttttagctcactaccctctgacttttcaaaggccgattcatataactggcagtcgagtt cacactgtaacagtgagtgtagcgcaaaaggtgcaagctactttgccctttataatcattc agaatgttattgtggtgatactaatccatctggttcggaatctacttcctcttcatgtaatacg tattgctttggttacagcagtgagatgtgtggtggtgaagatgcctattctgtgtaccaactt gactctgacacaaatagcaatagcataagcagctcagattcaagtacggagtctacttc tgcgtcgtcttccacaacttcttcaacaacgtcctccacaacatcaactacatcatcgact acatcatcaactacatcatcaatggcgtcttcctctacagtacagaattcccccgagtca actcaagcagctgcctctatttcaacgtcacagagttcgagcactgtaacgtcagaaag ctcgctaacttcggatactttggcaacgagcagtactagctctcaatcacaggacgcca cttcgataatctattccactacttttcacactgagggcggttccacaatttttgtcacgaaca ccatcacggcaagtgcacagaattcaggatcggccacaggtacagctggttctgattct acttctggcagcaaaacccacaaaaagaaggccaatgtaggggcaatcgtgggcg gtgtagtagggggtgtggttggagcggtagctattgcgttgtgcatacttcttatagtaagg cacatcaatatgaaacgtgaacaagaccgtatggagaaggaagcccaggaggcga taaagccggtggaatatcccgataagctatatgcagacgattttgatgataaccatggc ccgtctagtggatcattcgaggaagaacacactaaggggcagactgacatagctgca atagatgattctcgtcgtatatctaatgggacatttatcaacggtgggccaggtggcaag aataatgtcgatctagaggggagtacaatgtcaggg |
| pFixK | gtcgtggcagctaagtaatttcccttagtgatctaacccaatttctccaaatcacgcgagtt ggcaatctgtccccacatcgaacggggaataattcggcatatatatatgtgcgcgtatat acatgattatatggcatgtatgtgctctgtatgtatataaaactcttttttctttttttctctaaattt tttttccttatacattaggacctttgcagcataaattactatacttctata |
| sFixL | atgttatcatttacaaccaaaaattcctttaggttacttctattgattctgagttgcatttccacc attagggcgcaatttttttgttcaaagtagctcctctaactcaagcgcggtctcaaacctggt tgcacagcttgaaaacgaagtcgcatccctggaaaacgagaatgaaacgctgaaaa aaaagaacttgcacaagaaagacttaattgcctatttggaaaaggagattgctaaccta agaaaaaaaatagaagaaggttctgattctacttctggcagcaagttagctaagataat cacaggtactgtcatcgccattggagtatttgtaattctacttacgctgccactagctcact gggccgtgcagcccatcgtcagattacaaaaggctacggagttaattaccgagggca ggggattacgtccctccacaccgaggacgatatctagggcatccagtttcaaaagagg cttctcaagcggtttcgcagtcccatcttcccttttacaattcaacacagcggaggctggat caactacgtccgtaagcggtcacggagggtccgggcatggtagtggtgccgctttttca gctaatagctctatgaaaagcgcgatcaatcttggaaatgagaaaatgagtcctcctga ggaagagaacaaaatccccaacaatcataccgacgcgaaaatctcaatggatggct cattaaatcatgacttgttgggtccccacagtcttaggcataacgacactgatagaagttc taataggagtcacattttaacgacttcagccaatctgacggaagctcgtcttcccgactat cgtcgtttattttcagatgagttgagtgacctgacggaaaccttcaacacaatgacggac gctttggatcaacattacgccttgctagaggagagggtaagggcaaggactaaacaa ctggaggcggcaaagatagaggcagaagctgctaatgaggcgaagactgtgttcatt gcaaatatctcccatgaacttaaccagccattgtccgccatcgcaaactattctcacggtt gtacacgtttattaagggacatggacgacgcagtcgcgacacgtataagagaagccc ttgaggaggtcgcgagtcaaagtctacgtgctggacaaattattaaacacctgagaga gttcgttacgaaaggcgagaccgagaaagctccggaagatatacgtaaattagtgga ggagagcgcggcattagccctagttgggagcagagagcaggggggtccgtaccgtgtt cgagtatttgcccggtgcagagatggtcttggtcgaccgtatccaagtacagcaagtact aataaatcttatgcgtaatgcaatagaagccatgagacatgtagacaggagggaact aactatacgtactatgcctgctgatccgggcgaagtcgcggtggtcgttgaggacaccg |

| | |
|---|---|
| | ggggtgggatacctgaggaagtcgccggtcagctatttaagcccttcgttaccactaag<br>gctagtgggatggggattggactttctatcagtaaacgtatcgtcgaggctcatggaggc<br>gagatgacagtctctaaaaatgaggcaggaggagcaacatttaggtttacacttcctgc<br>atatcttgatgagaggatagtagctaatgac |
| VP64-FixJ | atggatgcgttggacgatttcgatcttgacatgttagggagtgatgcattggatgatttcga<br>tttagatatgttaggaagcgacgccttagacgacttcgatcttgatatgcttggctcagatg<br>ctttagacgattttgacctggatatgttaataaaccgtgcggatgggagtgggagatcag<br>gagttgatggaggcggttctatgactgattacacggtacatatcgttgatgacgaagagc<br>cagttaggaagtccctagcctttatgctaacaatgaatggcttcgcagtcaagatgcacc<br>agtcagcagaggcatttcttgctttcgcccccgatgtaagaaatggtgtactggttacgg<br>atctgaggatgccggatatgagtggagtagagctacttagaaacctgggcgacttaaa<br>aatcaatatcccgagcattgtcatcaccggccacggagacgttcccatggccgtggaa<br>gcaatgaaagcgggggcggttgactttatagaaaaaccgttcgaggatacggtcataa<br>tagaggccatcgaaagggcctcagaacaccttgtggctgcggaggcggacgtagac<br>gacgcgaacgacatcagggcaaggttgcagacgttgagtgagagagaaaggcagg<br>tgctgtcagctgtagttgcggggcttccaaataaaagtatcgcatatgatcttgacattag<br>cccacgtactgtagaggttcaccgtgctaatgtaatggcaaagatgaaagccaaaagt<br>cttccacacttagtgcgtatggcgcttgcaggaggatttggtccgagttaa |
| VP64-$FixJ_{nd}$ | atggatgcgttggacgatttcgatcttgacatgttagggagtgatgcattggatgatttcga<br>tttagatatgttaggaagcgacgccttagacgacttcgatcttgatatgcttggctcagatg<br>ctttagacgattttgacctggatatgttaataaaccgtgcggatgggagtgggagatcag<br>gagttgatggaggcggttctatgacagattacaccgtacacatcgtagatgacgagga<br>gccggtgcgtaaatcacttgcatttatgcttaccatgaacggatttgcggtaaagatgca<br>ccaatctgccgaggctttccttgcgtttgcgccagacgtgaggaatggggtgttagttaca<br>caattaaggatgcccgacatgtctggcgtggaattgctaagaaatcttggagatttgaaa<br>atcaatatcccatccatcgtgattacaggacatggagacgtgccgatggccgtagaag<br>ctatgaaggctggagccgttgatttcattgaaaaaccgtttgaggatacagtaattattga<br>ggctatagagagagcgagcgaacacttagtcgctgccgaggctgatgtcgatgatgc<br>caatgacatcagagctagattacaaactctttccgaaagggaaaggcaagtattgtctg<br>ccgtggtggccggactgccaaacaagtctatagcttatgatcttgacatttctcctaggac<br>tgtagaggttcaccgtgcgaacgtcatggcgaaaatgaaagccaagtccttgccccatt<br>tggtcaggatggccctagcaggtggctttggaccgagttaa |

**Table S3**. **Yeast strains summary**

| Strain name | Composition | Source |
|---|---|---|
| CEN.PK2-1C | MATa; his3D1; leu2-3_112; ura3-52; trp1-289; MAL2-8c; SUC2 | Kind gift form Dr. Luis Rubio |
| BY4741 | S288C isogenic yeast strain: MATa; his3D1; leu2D0; met15D0; ura3D0 | Kind gift form Dr. Luis Rubio |
| 'Kin' | CEN.PK2-1C<br>eLeu; eUra; dNeon reporter | This study |
| 'Intruder' | BY4741<br>eLeu; eUra; eHis | This study |
| 'RLM1' | CEN.PK2-1C<br>eLeu; eUra; RLM1 reporter | This study |
| 'STL1' | CEN.PK2-1C<br>eLeu; eUra; STL1 reporter | This study |
| 'OCH1' | CEN.PK2-1C<br>eLeu; eUra; OCH1 reporter | This study |
| 'synSSRE' | CEN.PK2-1C<br>eLeu; eUra; synSSRE reporter | This study |
| 'Kin synZIP' | CEN.PK2-1C<br>ZIPsln(SynZIP2); ZIPsln(SynZIP1); dNeon reporter | This study |
| 'Kin Δflo11' | CEN.PK2-1C Δflo11<br><br>eLeu; eUra; dNeon reporter | This study |
| 'Kin FLO11OX' | CEN.PK2-1C<br>eLeu; pGAL1::FLO11; dNeon reporter | This study |
| 'Intruder FLO11OX' | BY4741<br>eLeu; pGAL1::FLO11; eHis | This study |
| 'Kin sFLO11' | CEN.PK2-1C<br>eLeu; sFLO11; dNeon reporter | This study |
| 'Intruder sFLO11' | BY4741<br>eLeu; sFLO11; eHis | This study |

| | | |
|---|---|---|
| 'Intruder dNeon' | BY4741<br>eLeu; eUra; dNeon reporter | This study |
| 'Intruder dNeon FLO11OX' | BY4741<br>eLeu; pGAL1::FLO11; dNeon reporter | This study |
| 'FixJ/FixL' | CEN.PK2-1C<br>FixJ; sFixL; pFixK | This study |
| 'FixJ$_{nd}$/FixL' | CEN.PK2-1C<br>FixJ$_{nd}$; sFixL; pFixK | This study |
| 'FixJ free' | CEN.PK2-1C<br>FixJ; eUra; pFixK | This study |

**Video S1.** Time-lapse imaging of ‘Kin’ and ‘Intruder’ growth competition in the microfluidic device**.** ‘Kin’ is expression dNeon reporter (in cyan), ‘Intruder’ is marked by constitutively expressed mCherry (in red). Pictures were taken every 10 min from 15h up to 40h. Color coding as in Fig. 2E.
